# Successful dendritic cell vaccines require lasting in-situ TNFa secretion to license antitumor CD8^+^ T cell cytotoxicity

**DOI:** 10.64898/2026.04.06.716539

**Authors:** Nadine Santana-Magal, Aseel Khateeb, Kfir Inbal, Annette Gleiberman, Ayelet Kaminitz, Tomer Weiss, Gal Verbin, Alon Richter, Amichai Zarfin, Leen Farhat-Younis, Amit Gutwillig, Adi Frish, Eric Shifrut, Inbal Reuveni, Adi Barzel, Carmit Levi, Peleg Rider, Matthew H. Spitzer, Edgar G. Engleman, Asaf Madi, Yaron Carmi

**Author notes:** Corresponding authors: Yaron Carmi and Asaf Madi The Department of Pathology, Gray School of Medicine, Tel Aviv University, Tel Aviv 69978, Israel. Asterisk denotes coequal contribution (Nadine Santana-Magal, Aseel Khateeb, Kfir Inbal). Grant Support: This work was funded by the Fritz Thyssen Foundation, the Israel Cancer Research Foundation, and the Israel Science Foundation (ISF) (Grant Number 2262/18).

## Abstract

Dendritic cells (DCs) are central to activating cytotoxic CD8⁺ T cells, yet DC-based vaccines have achieved limited success against established tumors. To address this gap, we analyzed the transcriptomic and functional changes CD8⁺ T cells undergo following interactions with DC subsets in lymphoid organs and tumor sites. This approach allowed us to map their trajectory from naïve to fully cytotoxic effector cells. We found that classical DCs in lymphoid organs provide essential antigen presentation but fail to elicit cytotoxicity. Instead, antigenexperienced CD8⁺ T cells require additional inflammatory signals, primarily through TNFα, delivered at tumor sites by infiltrating myeloid DCs. Effective cytotoxic responses therefore depend on the synchronization of these distinct, temporally separated signals. Notably, tumor antigen–pulsed DC vaccines rapidly lose TNFα expression after infiltrating tumors, limiting their efficacy. These findings establish a sequential model of T cell activation and suggest strategies to enhance the potency of DC-based immunotherapies.

## Introduction

Since the discovery of dendritic cells (DC) by Ralph M. Steinman and Zanvil A. Cohn, countless studies have highlighted their central role in determining T cell tolerance or activation towards specific antigens, including in cancer. Therefore, deciphering the types of T cell-DC interactions that either promote activation or tolerance, and the receptors/ligands that mediate them, has been the source of extensive research lasting over several decades^2^. DC are composed of heterogeneous cell populations that differ from each other in their pattern-recognition receptors, tissue distribution, and migratory and antigen presentation capabilities ^3,4^. Based on our current understanding of DC ontogeny, they are comprised of plasmacytoid DC (pDC), classical DC type 1 (cDC1), classical DC 2 (cDC2) and monocyte-derived DC (MoDC)^3,5^. Lymph node-resident or migratory cDC1 and cDC2 are considered as the main antigen presenting cells and are equipped with the ability to migrate to the T cell zone. cDC2s are thought to predominantly present MHC II-associated antigens to CD4^+^ T cells, while cDC1s specialize in presenting antigens to CD8^+^ T cells. Yet, cDC1s have a unique ability to crosspresent exogenously acquired antigens to both CD4^+^ and CD8^+^ T cells, and mice deficient in cDC1s fail to mount T cell immunity against tumor antigens^1,6^

The remarkable capacity of DCs to activate cytotoxic T cells has made them attractive targets for tumor vaccines. Although cDC1 subsets are superior antigen-presenting cells and promote extravasation of T cells into tumor, their prevalence in tumors and blood is limited^7–10^. In contrast, MoDC can be generated easily from circulating or tumor-infiltrating monocytes ^11^; thus, most clinical trials take advantage of their relative prevalence ^12,13^.

Traditional efforts focused on finding the most potent stimuli to mature and activate DC and on identifying the ideal, unique, high-avidity tumor antigens that can be loaded on the DC ^14,15^. Vaccination with neoantigens loaded on activated DC, or injected with TLR3 agonists, elicits strong T-cell immunity and prevents tumor recurrence in patients with resected melanoma ^16,18^. Nonetheless, attempts to employ DC to treat established tumors, rather than as prophylactic tumor vaccines, have achieved only marginal success in clinical trials ^19^. In melanoma, for example, clinical trials with autologous DC vaccines have been shown to induce tumor-specific immune responses in both lymph nodes and metastases. These studies include adjuvant-based DC vaccines ^15^, mRNA transfection of monocyte-derived DC^20^ or *ex-vivo* generated DC loaded via nanoparticles^21^. Despite induction of reactive CD8+ T cells, objective clinical responses were rarely observed. Interestingly however, vaccination of high-risk melanoma patients with peptides in combination with Poly:ICLC or mRNA encoding tumor neoantigens after surgical resection resulted in significant reduction in tumor recurrence^17,18,22^. While the presence of a suppressive microenvironment in established tumors is likely to inhibit tumor-reactive T cells, clinical data indicate that tumor-reactive T cells do infiltrate tumors following vaccination ^23,24^. On one hand, clinical outcomes highlight the potential importance of DC- vaccines but also emphasize the current limitations of harnessing DC against established cancers. Therefore, why DC vaccines fail to treat tumors despite their remarkable potential to elicit T-cell immunity is not clear.

To study this gap, we performed an in-depth analysis of the transcriptomic and functional changes that CD8^+^ T cells undergo following interactions with DC, resulting in a detailed description of their trajectory from naïve to fully cytotoxic effector cells. We found that antigen presentation to CD8^+^T cells is mediated predominantly by classical DC (cDC) in lymphoid organs, yet it is not sufficient to elicit T cell cytotoxicity. Instead, antigen-experienced T cells must receive an additional licensing signal from myeloid dendritic cells (MoDC) in the tumor microenvironment. Synchronization of these two activation modules is requisite for eliciting T cell cytotoxicity. Overall, this study provides new insights into spatio-temporal DC organization and the signals that are required to elicit T-cell immunity and a framework to increase the potency of DC vaccines.

## Materials and Methods

### Mice

Wild-type (WT) C57BL/6 mice were obtained from Envigo (Jerusalem, Israel). B6;129SGt(ROSA)26Sortm1.1(CAG-COX8A/Dendra2)Dcc/J mice, which express Dendra2 green/red photoswitchable monomeric fluorescent protein in mitochondria were purchased from Jackson Laboratories (Bar-Harbor, ME, USA). B6.Cg-Ptprca Tg (UBC-PA-GFP)1Mnz/J mice, which express constitutively photoactivable GFP were obtained from Dr. Ziv Shulman at the Weizmann Institute, Israel. Gp100 mice were obtained from Jackson Laboratory. OT1 mice were also obtained from Jackson Laboratory. All mice were housed in an American Association for the Accreditation of Laboratory Animal Care–accredited animal facility and maintained under specific pathogen-free conditions. Animal experiments were approved and conducted in accordance with Tel-Aviv University Laboratory Accreditation #01-16-095. Male and female 8-12 weeks old mice were used in all experiments.

### Cell lines

B16F10 cells (CRL-6475) were purchased from ATCC, in January 2017, and *Ret* melanoma cells were a kind gift from Prof. Neta Erez at Tel Aviv University. A375 and YUMM1.7 cell line was a kind gift from Prof. Tamar Geiger at Tel Aviv University. HEK-293FT were purchased from ThermoFisher Scientific (Waltham, MA) in January 2017. Cells were cultured in DMEM supplemented with 10% heat-inactivated FBS, 2 mM L-glutamine, and 100 µg/mL penicillin/streptomycin under standard conditions (all from Biological Industries, Israel). Melan-A cells were purchased from Welcome Trust Functional Genomics Cell Bank, London, UK in February 2019. Cells were cultured as described above with the addition of 200 nM of Phorbol 12-myristate 13-acetate (PMA) (Santa-Cruz biotechnology, Ca). YUMM 1.7 were cultured in DMEM:F12 supplemented with 10% heat-inactivated FBS, 2 mM L-glutamine, 100 µg/mL penicillin/streptomycin and 1% MEM-Eagle non-essential amino acids. All cells were routinely tested for mycoplasma using an EZ-PCR Mycoplasma Test Kit (Biological Industries, Israel) according to the manufacturer’s instructions.

### In vivo tumor models

For melanoma tumor studies, 2×10^5^ B16F10, 2×10^5^ YUMM1.7, or 5×10^5^ *ret* cells were suspended in 50 µL of appropriate medium and were injected s.c. to C57BL/6 mice above the right flank, and the size of growing tumors was measured twice a week using calipers. Treatment was applied at different days post-injection. When tumors reached 200 mm^2^, the mice were sacrificed due to ethical considerations.

### DC isolation

All tissue preparations were performed simultaneously from each individual mouse (after euthanasia by CO_2_ inhalation). Peripheral blood was collected through perfusion, where the right atrium of the heart was cut and HBSS supplemented with 10 mM EDTA was circulated to the left ventricle of the heart and blood was collected into a sodium heparin-coated vacuum tube. In a laminar hood, the blood was pipetted on a Histopaque-1077 Hybri-Max (Sigma Aldrich, Merck, Israel) density gradient medium and centrifuged at 400 rcf for 15 min at room temperature with no brakes. Mononuclear cells were collected into a 15 mL tube and dissolved in 50 mL of complete RPMI 1640, washed (400 rcf for 5-10 min at 4⁰C) and resuspended in 500 µL complete RPMI 1640.

Spleen and LN were homogenized in HBSS supplemented with 2% FBS and 5 mM EDTA at 4⁰C and strained with 70 µM strainer (ThermoFisher Scientific, Waltham, MA). The cells were centrifuged and resuspended in 500 µL complete RPMI 1640. Tumors from human or mice were digested in RPMI 1640 with 2 mg/mL collagenase IV, 2,000 U/mL DNase I (both from Sigma Aldrich, Merck, Israel), for 40 min at 37°C with magnetic stirrers (200 rpm). Cells were homogenized in RPMI 1640 with 10% FBS and strained in 70 µM cell strainer (ThermoFisher Scientific, Waltham, MA). Cells were then washed and resuspended in 500 µL complete RPMI. Utilizing magnetic CD11b MicroBeads and column (both from Miltenyi Biotec), CD11b^+^ positive cells were isolated from tumor lysate, PB, and DLN according to the manufacturer’s instruction. Isolated cells were let to adhere for 2 hours in complete RPMI 1640 in 37°C and loosely adhered cells were further sorted by FACSAriaII as FSC^low^SSC ^low^ cells.

Bone marrow cells were isolated from the Tibia, Femur and Hip bones of control C57Bl/6 mice under sterile conditions in a laminar hood. Bones were washed extensively in PBS (Gibco), incubated for 20 sec in 70% EtOH and washed again twice in PBS. Bones were grind using sterile Mortar and Pestle and filtered through 70 uM strainer(ThermoFisher Scientific). Cells were washed and re-filtered and cultured in 15 cm culture dishes for 4-5 days in complete DMEM (GIBCO) the presence of 50 ng/ml GM-CSF and 10 ng/mL IL-4 (PeproTech, Rocky Hill, NJ) to generate monocytes-derived DC.

### T cell isolation

All tissue preparations were performed simultaneously from each individual mouse (after euthanasia, by CO_2_ inhalation). For isolation of T cells from the spleen and lymph node, the organs were removed from euthanized mice and mashed through a 70 µM cell strainer. Cells were then washed by centrifugation at 600 rcf for 5 min. at 4-8°C. For isolation of tumorinfiltrating T cells, tumors were enzymatically digested with 2 mg/mL collagenase IV and 2,000 U/mL DNase I (both from Sigma Aldrich, Merck, Israel) in HBSS for 30 min at 37°C with magnetic stirrer (200 rpm). Cells were then washed by centrifugation at 600 rcf for 5 min. at 4-8°C.

For T cells from peripheral blood, peripheral blood was collected with perfusion of the animal and transferred into sodium heparin-coated vacuum tubes before to 1:1 dilution in HBSS supplemented with 2% FBS and 5 mM EDTA. Lymphocytes from peripheral blood and spleen were enriched on a Histopaque-1077 Hybri-Max (Sigma Aldrich, Merck, Israel) density gradient medium and collected PBMC were washed twice with a complete RPMI 1640. For all tissues, cells were then incubated with anti-CD4 or anti-CD8 magnetic beads (MojoSort™ Nanobeads, BioLegend, Carlsbad, CA) according to the manufacturer’s instruction.

### T-cell culture and expansion

T cells were cultured in RPMI 1640 supplemented with 1% Pen-Strep, 10% heat-inactivated FBS, 1% Sodium pyruvate, 1% MEM-Eagle non-essential amino acids, 1% InsulinTransferrin-Selenium (PeproTech, Rocky Hill, NJ), 50 µM β-mercaptoethanol (SigmaAldrich, Merck, Israel) and were supplemented with 1,000 IU/mL recombinant murine IL-2 (PeproTech, Rocky Hill, NJ) on tissue culture plates pre-coated with 0.5 µg/mL anti-CD3 antibodies (clone 17A2, Biolegend).

### Adoptive T Cell Transfer

For immunotherapy assays using adoptive transfer of tumor-infiltrating T cells, mice were injected s.c. with B16F10 and tumors were let to grow for 8-10 days, until they reached an average size of 25 mm^2^. 1×10^6^ of splenic CD8^+^ T cells infected with TCR recognizing gp100 ^25^ per mouse were injected i.v. into tumor-bearing mice sublethally-irradiated with a single dose of 600 rad, followed by i.p. injections of 300,000 IU of IL-2 (PeproTech, Rocky Hill, NJ) for four consecutive days. To increase DC viability in late-stage tumors, non-irradiated recipient mice were injected i.p with 80 µg/mouse anti-CD40, and with 1 µg/mouse of recombinant GM-CSF prior to CD8^+^ transfer.

### In vitro Killing assay-

Mouse CD8^+^ T cells were isolated from various organs of naïve or tumor-bearing mice. CD8^+^ T cells were co-cultured with H2B-Tdt labeled tumor cells (5,000 cells per well) with or without CD11b^+^ DC at a ratio of 1:5:2 (T:E:DC), or with CD11b DCs’ supernatant. For human, CD8 T cells were isolated from propherial blood, infected with 1G4-TCR retrovirus, cells were co cultured with H2B-tdt A375 melanoma cells (10,000 cells per well) with or without LPS activated MoDCs’ supernatant.

Assays were conducted in a flat bottom 96-well plate. Cells were imaged by IncuCyte S3 /S5 imager (Sartorius), and images were then used to calculate numbers of target cells or confluency by IncuCyte software (version 2021A/2022B, basic analyzer). The number of melanoma cells was normalized to the number of melanoma cells in each well at time 0.

### DC vaccines

For prophylactic vaccination, classical splenic cDCs and MoDCs cultured from blood monocytes were isolated on CD11c and CD11b magnetic beads (Miltenyi Biotec), respectively from C57Bl/6 mice. Cells were then activated overnight with 1 µg/mL of poly I:C or LPS (InvivoGen) and pulsed with 5 µM of MHC-I-restricted peptides gp100 (EGSRNQDWL) and TRP1 (TAPDNLGYM) (Syntezza Bioscience) for an additional one hours. Peptide-loaded DCs were washed, resuspended in RPMI, and injected s.c three consecutive times, five days apart. at concentration of 2 × 10⁶ cells per 100 µL. Three days after the third vaccination, mice were challenged subcutaneously in the right flank with 2 × 10^5^ B16F10 melanoma cells. For vaccination of tumor-bearing mice, C57Bl/6 mice bearing day six B16F10 tumors were injected s.c twice four days apart with LPS-activated DC pulsed with the above melanoma antigens.

### Adoptive T Cell Transfer and Tumor Recurrence Model

For adoptive T cell transfer ACT, two weeks after the second vaccination, mice were euthanized, and CD8⁺ T cells were isolated from spleens on a Ficoll gradient (Sigma) followed by negative selection using magnetic beads (Miltenyi Biotec). T cells were incubated at 37oC to restore homeostasis and 5×10⁶ cells per mouse were injected into the tail vain of sub-lethally irradiated C57Bl/6 mice. After 3 days, mice were challenged subcutaneously with 2×10^5^ B16F10 melanoma cells and tumor growth was measured by caliper twice a week.

For tumor recurrence study, mice were challenged with 2×10^5^ B16F10 melanoma cells s.c. in the right flank, and tumors were allowed to grow until they reached an average size of approximately 100 mm². Mice were then anesthetized with i.p. injection of ketamine (100 mg/kg) and xylazine (4 mg/kg), and the surgical site was sterilized with 70% ethanol. A small incision was made to carefully expose and excise the tumor while minimizing damage to surrounding tissues, after which the incision was closed with wound clips. Postoperative care included administration of analgesics and monitoring for signs of distress or infection. Mice were allowed to recover for one week before receiving immunotherapy. Peptide-loaded cDCs and MoDCs were prepared as described above and administered s.c. at a dose of 2×10⁶ cells per mouse to the corresponding treatment groups.

### Dendritic Cell Trafficking to Tumors

To assess DC infiltration into tumors, splenic cDCs and MoDCs pulsed with MHC-I-restricted melanoma antigens gp100 and TRP1 were prepared from CD45.1 mice and injected into B16F10 tumor-bearing CD45.2 mice, when tumors reached an average size of 50 mm². Fortyeight hours post-vaccination, tumors were harvested and analyzed by flow cytometry for the presence of MHC-II⁺CD11c⁺CD45.1⁺ cells, indicating DC migration to the tumor microenvironment.

### Lentiviral infection

For preparation of lentivirus, 1.3×10^6^ HEK-293FT cells were plated on a 6-well plate precoated with 200 µg/mL poly-L-Lysine (Sigma Aldrich, Merck, Israel) and left to adhere overnight. pLVX plasmids containing wasabi, or tdTomato under an EF1 promoter (kindly provided by Dr. Rathicker-Flynn, Stanford University) were mixed with psPAX2 (a gift from Didier Trono, Addgene plasmid # 12260) and pCMV-VSV-G (a gift from Bob Weinberg, Addgene plasmid # 8454) at a molar ratio of 3:2:1 and cells were transfected using Polyplus jetPRIME® reagent (Polyplus transfections). After 24 hours, media was replaced with complete DMEM supplemented with 0.075% Sodium Bicarbonate. Media-containing viruses were collected after 24 h and 48 h. For infection, virus-containing media were mixed with 100 ug/mL polybrene (Sigma Aldrich, Merck, Israel), added to B16F10 cells for 30 min at 37°C 5% CO2, and then cells were centrifuged for 30 min. at 37°C and 500 rcf. Medium was replaced with complete DMEM supplemented with 0.075% Sodium Bicarbonate. After three days in culture, cells that expressed wasabi, TRP1-GFP, or tdTomato were sorted by FACSAriaII and were tested for mycoplasma contamination by EZ-PCR Mycoplasma Test Kit (Biological Industries, Israel).

### Retroviral infection

For human retroviral preparation, 1×10^6^ HEK-GP2 cells were plated on a 6-well plate precoated with 200 µg/mL poly-L-Lysine (Sigma Aldrich, Merck, Israel) and left to adhere overnight. Medium was replaced to media without antibiotics and then cells were cotransfected with a 2:1 molar ratio of PMSGV1 and RD114 plasmids (both were kindly provided by Prof. Dario A. A. Vignali) using lipofectamine 2000 reagent. After 24 hours, the media was replaced with complete DMEM supplemented with 1mM Sodium Bicarbonate. Media-containing viruses were collected after 24 hours and 48 hours. Prior to infection, CD8 T cells were cultured on a plate pre-coated with anti-CD3 (0.5 µg/mL) and anti-CD28 (0.2ug/ml) in T cell media containing high-dose IL-2 (500 IU/mL) for 2 days. Next, 1 mL of retroviruses per well was centrifuged for 2h at 4000rpm at 32°C. Then virus media was replaced by 1ml of 3×10^5 CD8^+^ T cells. After additional centrifugation for 10 min with acceleration and brakes 1, T cells were cultured for additional three days in T cell media containing high-dose IL-2 (1000 IU/mL).

### Mice Data: Bulk RNA-seq Sequencing

Library preparation, sequencing, and primary data processing (adapter/Poly-A trimming and alignment) were performed by the following core facilities of the Nancy and Stephen Grand Israel National Center for Personalized Medicine, Weizmann Institute of Science: The Mantoux Bioinformatics Institute; The Crown Genomics Institute; The De Botton Protein Profiling Institute; The Wohl Drug Discovery Institute; and The Medicinal Chemistry Institute. 44 samples were received for library prep, sequencing, and analysis. Libraries were prepared using the INCPM mRNAseq protocol. 75bp single reads were sequenced on a NextSeq High output. Sequencing Libraries were prepared using Bulk MARS-Seq. 75 SR reads were sequenced on 1 lane(s) of an Illumina NextSeq High output. The output was ∼10 million reads per sample.

### Mice Data: Bulk RNA-seq Analysis

Poly-A/T stretches and Illumina adapters were trimmed from the reads using cutadapt [<u>6</u>]; resulting reads shorter than 30bp were discarded. Remaining reads were mapped onto 3’ UTR regions (1000 bases) of the M. musculus, GRCm39 genome according to Ensembl annotations, using STAR [<u>2</u>] with EndToEnd option and outFilterMismatchNoverLmax was set to 0.05. Deduplication was carried out by flagging all reads that were mapped to the same gene and had the same UMI. Counts for each gene were quantified using htseq-count [1,<u>3</u>], using the gtf above. UMI counts were corrected for saturation by considering the expected number of unique elements when sampling without replacement. Gene counts were imported into R (v4.3.2), and differential expression was performed with DESeq2 (v1.42.1). Ensembl gene IDs were annotated to MGI symbols via the “mmusculus_gene_ensembl” dataset to standardize nomenclature. Genes with fewer than ten reads across all samples were excluded. Counts were normalized using the regularized log (rlog) transformation. Principal component analysis identified one MoDC control outlier (Naive_MoDC_M4), which was removed.

Pairwise contrasts (treatment vs. control) employed the Wald test; multi-group comparisons (cDC_6H, cDC_Naive, MoDC_6H, MoDC_Naive) used the likelihood ratio test (LRT) to assess overall group effects. Differentially expressed genes (adjusted p-value < 0.05) were visualized in row-wise Z-scored heatmaps, with ligand–receptor modules curated from NicheNet.

### Mice Data: Algorithm for DSA Calculation and Ligand–Receptor Interactions

Canonical ligand-receptor pairs were obtained from NicheNet[36]. Circos plots (circlize v0.4.16) illustrated interactions between activated MoDC ligands and CD8⁺ T cell receptors. For downstream activation (DSA) scoring, we adapted the InterFLOW [37] algorithm developed by the Madi Lab. The following steps outline the computational process:

1. **Data Preparation:** rlog-normalized CD8⁺ T cell counts (DESeq2 rlog) were min-max scaled; mean gene expression per gene was computed.
2. **Correlation Matrix:** A Pearson correlation matrix was generated; absolute values of negative correlations were used. Interactions were filtered against a PPI table from Mippie v1.0 [38].
3. **TF Enrichment:** Hypergeometric tests (SciPy’s hypergeom.sf) identified enriched TFs (FDR < 0.05).
4. **Network Construction:** An undirected graph G=(V,E) was built: enriched TFs connected to a virtual sink (infinite capacity edges), other edges weighted by (mean expression × correlation). TF-target mappings used CellTalkDB [39].
5. **Max Flow Calculation:** The Dinitz algorithm from NetworkX (v2.8.4) [40] computed max flow from each receptor to the sink.
6. **Significance Testing:** A 200-iteration permutation test shuffled target genes; observed flows were compared to the null via one-sample t-tests (α=0.05).

### Mice Data: Proteomics Analysis

BMDCs were isolated from the bone marrow of mice and cultured for one week with 50 ng/mL GM-CSF to differentiate into MoDCs. The day before LPS activation, the culture medium was replaced with one lacking FBS. Half of the cells were activated with LPS for 24 hours, while the remaining cells were left unactivated. After 24 hours, the supernatant was collected and purified using an Amicon Ultra Centrifugal Filter (MWCO 30 kDa), suitable for protein purification. In total, six MoDCs supernatant samples (three activated with LPS and three naïve) were sent for proteomics analysis at Smoler Proteomics Center, Technion.

- **Proteolysis and Mass Spectrometry Analysis** The protein samples were brought to 8.5M Urea, 100mM ammonium bicarbonate and 10mM DTT. Protein amount was estimated using Bradford readings. The samples were reduced (60°C for 30 min), modified with 35.2mM iodoacetamide in 100mM ammonium bicarbonate (room temperature for 30 min in the dark) and digested in 1.5M Urea, 17.6mM ammonium bicarbonate with modified trypsin (Promega), overnight at 37°C in a 1:50 (M/M) enzyme-to-substrate ratio. An additional second digestion with Trypsin was done for 4 hours at 37°C in a 1:100 (M/M) enzyme-tosubstrate ratio.

The resulted peptides were analyzed by LC-MS/MS using the Exploris 480 mass spectrometer (Thermo) fitted with a capillary HPLC (EV-1000, Evosep (480-Ex1)). The peptides were loaded onto a 15cm, ID 150µm, 1.9-micron Endurancse column EV1137 (Evosep). The peptides were eluted with the built-in Xcalibur 15 SPD (88 min) method.

Mass spectrometry was performed in a positive mode using repetitively full MS scan (m/z 350–1200) followed by High energy Collision Dissociation (HCD) of the 20 most dominant ions (>1 charges) selected from the full MS scan. A dynamic exclusion list was enabled with exclusion duration of 30s

- **Data Processing and Statistical Analysis** The mass spectrometry data was analyzed using the MaxQuant software version 2.1.1.0 [^41^] for peak picking and identification using the Andromeda search engine, searching against the Mus musculus proteome from the Uniprot database (October 2024) with mass tolerance of 6 ppm for the precursor masses and 20 ppm for the fragment ions. Oxidation on methionine and protein N-terminus acetylation were accepted as variable modifications and carbamidomethyl on cysteine was accepted as static modifications. Minimal peptide length was set to seven amino acids and a maximum of two miscleavages was allowed. The data was quantified by label free analysis using the same software. Peptide- and protein-level false discovery rates (FDRs) were filtered to 1% using the target-decoy strategy. Protein table was filtered to eliminate the identifications from the reverse database and common contaminants. Statistical analysis of the identification and quantization results was done using Perseus 1.6.7.0 software [^42^]. Log2(0) values were replaced with 18.5, the threshold intensity. Proteins identified by a single unique peptide were not considered reliably identified.

The Smoler Center provided log₂ fold-change (log₂FC) and *p*-values (Student’s *t*-test) for each protein. For visualization, Python and Seaborn were used to generate volcano plots of log₂FC vs. -log₁₀(*p*). Proteins with |log₂FC| > 1 and *p* < 0.05 were considered significant. Key proteins (|log₂FC| > 2 and -log₁₀(*p*) > 5) were highlighted.

Gene Set Enrichment Analysis (GSEA) was conducted using the gseapy package in Python with the “KEGG_2019_Mouse” gene set. Genes were ranked using the metric –log₁₀(*p*) × sign(log₂FC) and analyzed via the preranked method. Pathways with FDR < 0.05 were considered significantly enriched. Top and bottom pathways were visualized in a scatter plot based on pathway size and NES. A GSEA enrichment plot was generated for the TNF signalling pathway using gseapy.plot.

### Human Data: CD8⁺ T Cells Single-Cell RNA-seq Analysis

Single-cell RNA-seq data were processed in R using Seurat [43]. Cells with < 200 or > 2,500 detected genes, total RNA counts outside 700–3,500, or > 3% mitochondrial UMIs were excluded. Remaining counts were log-normalized (NormalizeData) and scaled while regressing out mitochondrial percentage. Highly variable genes were identified by FindVariableFeatures. Dimensionality reduction employed PCA (RunPCA), with 12 PCs selected via ElbowPlot and FindPC. Neighbors and clusters were identified using FindNeighbors and FindClusters (resolution = 0.3), and visualization was achieved with UMAP (RunUMAP) and t-SNE (RunTSNE). Batch effects were corrected with Harmony (RunHarmony), after which PCA, UMAP, and t-SNE were repeated to confirm alignment. Marker genes per cluster were detected by FindAllMarkers (Wilcoxon test), with the top 20 for each cluster exported for annotation (Supplementary Excel 1). Feature plots were generated for *SELL*, *IL7R*, *CCR7*, *LEF1*, *MKI67* (naïve/proliferation), and *PRF1*, *CD44*, *CD69*, *HSPA1A*, *CXCL13*, *TNFRSF18*, *TYROBP* (cytotoxic/activated). Based on these markers, five effector CD8⁺ T cell clusters (Cytotoxic, Activated, Stress-Responsive, CXCL13⁺ T Helper–Like, Effector) were selected for further analysis. This subset was renormalized, variable features re-identified, and PCA/clustering repeated (elbow = 11). Differential expression between tumor and lymph node samples was performed with FindMarkers (adjusted p < 0.05) and visualized by DoHeatmap. A cytotoxic signature score (Supplementary Excel 2) was calculated with AddModuleScore, and violin plots comparing tumor vs. LN were generated; significance was assessed via Wilcoxon test in the ggsignif package.

### Human Data: All Cells Single-Cell RNA-seq Analysis

The full dataset underwent similar quality filtering (cells with < 100 or > 3,000 genes, total UMI 300–3,500, mitochondrial content > 4% removed). Data were log-normalized, scaled (mitochondrial regression), and the top variable features identified. PCA selected 13 PCs (ElbowPlot), followed by clustering at resolution = 0.3. Batch correction with Harmony was applied before repeating PCA, UMAP, and t-SNE; resulting t-SNE plots displayed 12 clusters and tissue distribution. A feature plot of *ITGAX* (CD11c) identified dendritic cell clusters (5 and 9), which were subsetted and re-analyzed (PCA elbow = 13, clustering as above). To assess enrichment of MoDC and cDC signatures in LN and tumor microenvironment, module scores were computed via AddModuleScore using gene sets in Supplementary Excel 2. Violin plots of enrichment scores were generated and statistical significance determined by Wilcoxon test with ggsignif.

### Immunohistochemistry

For frozen sections, mice and human tissues were fixed in 4% paraformaldehyde for 1 hour and equilibrated in a 20% sucrose solution overnight. Tissues were then embedded in frozen tissue matrix (Tissue-Plus O.C.T. Compound, Scigen) and frozen at - 80°C. The 5-µm-thick sections were blocked with 5% BSA and stained with 1:100 diluted primary antibodies. For anti-mouse staining, the antibodies used were MHCII (clone M5/114.15.2), CD11c (clone N418), CD11b (cloneM1/70), anti-TCRβ (H57- 597), all from BioLegend. In addition, Cleaved Caspase-3 (clone 5A1E) from Cell Signaling technologies and LYVE-1 (clone ALY7) from Invitrogen were used. Nuclei were counterstained with Hoechst 33342 (ThermoFisher Scientific, Waltham, MA). For anti-human staining, the antibodies used were HLA-DR (clone L243), CD11c (clone 3.9), both from BioLegend. Cleaved Caspase-3 (clone 5A1E) was also utilized. Microscopy was performed with a ZEISS LSM 800 confocal microscope and analyzed using ZEN software (ZEISS, Germany).

### Enzyme-Linked Immunosorbent Assay

Levels of secreted TNF-alpha cytokine in 24h LPS-activated, 48h after wash and LPS/CL307 reactivated supernatants of mouse/human MoDCs were quantified using a sandwich ELISA kit (R&D Systems, Cat. No. DY406). Briefly, 96-well plates were coated overnight at 4°C with capture antibody specific for TNF-alpha. After blocking with 1% BSA in PBS for 1 hour at room temperature, samples and standards were prepared and added to the wells and incubated for 2 hours. Following washes, biotinylated detection antibody was applied, followed by streptavidin-HRP and substrate solution (TMB). The reaction was stopped with Biofix stop solution (LSTP-1000-01) and absorbance was measured at 450 nm using a BioTek Synergy HT microplate reader. All samples were assayed in triplicates, and TNF alpha concentrations were determined by the standard curve equation.

### Confocal microscopy

CD11b^+^ DC isolated from tumor, DLN or PB were cultured on glass-bottom confocal plates (Cellvis, Mountain View, CA) in DMEM supplemented with 50 ng/mL GM-CSF. B16Wasabi, CD8^+^ T cells and CD11b^+^ DC were co-cultured on glass-bottom confocal plates in Tcell medium without IL-2 and incubated overnight under standard conditions. For functional assays, DC were applied with 1:50 diluted FITC-labeled latex beads (Sigma Aldrich, Merck, Israel) and incubated for 30 min before imaging. For visualization of DC activated with exosomes/lysosomes, DC were stained with LysoSensor (L7533, ThermoFisher Scientific, Waltham, MA). For visualization of lysosome trafficking, B16F10 cells were cultured on glass-bottom confocal plate with 1:1000 diluted lysotracker (L7526, ThermoFisher Scientific, Waltham, MA) for 30 min in 37°C and washed twice before the addition of 50 x 10^4^ MoDC isolated from C57BL/6 naïve mice. Images were collected using a Zeiss LSM800 confocal laser scanning microscope and analyzed using ZEN software (Carl Zeiss Microscopy).

### Flow cytometry

Cells were analyzed using flow cytometry (CytoFLEX, Beckman Coulter, Lakeview Indianapolis, IA) and sorted by FACS (BD FACSAria™ III, BD Biosciences, Franklin Lakes, NJ). Datasets were analyzed using FlowJo software (Tree Star). The following mice antigen were used: (Brilliant violet 421)-CD45 (clone 30-F11), (Alexa Fluor 647)-MHCII (clone M5/114.15.2), (Alexa Fluor 594, Brilliant violet 605)- CD11c (clone N418), (Per-CP)-CD11b (cloneM1/70), (Alexa Fluor 488)- CD86 (clone GL-1), (PE-Cy7)- CD103 (clone 2E7), (APCCy7)- Ly6C (clone HK1.4), (Brilliant violet 421)-βTCR (clone), (PE)- CD4 (clone RM4-4), (Brilliant violet 605)-CD8 (clone 53-6.7), (Alexa Fluor 488)- CD45 (clone 30-F11), (APC)NKp46 (clone 29AI.4), (PE/Cy7)- CD19 (clone 6D5), (PE)- B220 (clone RA3-6B2), (Alexa Fluor 488)- CCR7 (clone 4B12), (PE)-MERTK (clone 2B10C42).

For human specific antigens, the following antibodies were used: (Alexa Fluor 488) CD3 (HIT3a), (Alexa Fluor 594) CD4 (RPA-T4), (Allophycocyanin) CD19 (HIB19), (Alexa Fluor 647) CD8 (HIT8a), (Brilliant Violet 650) CD11c (3.9), (Alexa Fluor 647) CD16 (3G8), (PerCp/Cy5.5) CD32 (FUN-2), (Brilliant Violet 421) CD64 (10.1), (Allophycocyanin/Cy7) CD45RO (UCHL1), (Phycoerythrin/Cy7) CD45RA (HI100). All Abs were purchased from BioLegend. Cells were suspended in FACS buffer consisting of HBSS with 2% FCS and 0.05mM EDTA.

### Human Data: CyTOF Analysis

Mass cytometry (CyTOF) data were analyzed using FlowJo™ v10.8 software (BD Life Sciences) [44]. Analysis focused on the frequency and activation status of CD8⁺ T cells within the CD45⁺CD3⁺ gate. Specifically, the expression of CD69 and PD-1 on CD8⁺ T cells was quantified, and the expression levels of granzyme B (GZMB) were compared between tumor and matched lymph node (LN) samples. To assess statistical differences in GZMB expression between tumor and LN samples from the same patients, paired *t*-tests were performed in R using the t_test function from the rstatix package, with the paired parameter set to TRUE. The null hypothesis was that there was no difference in mean GZMB expression between tumor and LN-derived CD8⁺ T cells. A two-tailed *p*-value < 0.05 was considered statistically significant. Paired data visualization was performed using the ggpaired function from the ggplot2 package [45], generating boxplots with individual data points and annotated *p*-values to highlight differences in GZMB expression between tissue compartments.

### Statistical analyses

Each experiment was performed three times. Each experimental group consisted of at least three mice. For time course experiments, significance was calculated using the nonparametric two-way ANOVA with Tukey’s correction for multiple hypotheses. In some cases, Bonferroni-Sidak post-test was performed after two-way ANOVA. For two groups analysis, one-way ANOVA with Dunn’s test was performed. The results were analyzed by Prism (GraphPad Software, Inc.). All statistical analyses were performed in Prism (GraphPad Software, Inc.)

### Study approval

All animal protocols were approved by the Stanford University Institutional Animal Care and Use Committee under protocol: 01-16-095. The Tel Aviv University Institutional Review Board approved the human subject protocols, and informed consent was obtained from all subjects prior to participation in the study.

## Results

### DC vaccines fail to eradicate established tumors despite induction of tumor-reactive T cell clones in the draining lymph nodes

To study what factors limit the therapeutic efficacy of DC vaccines, we established several cancer mouse models aimed to recapitulate the clinical settings and outcomes of DC vaccines. Initially, DC capacity to promote tumor-protective immunity was tested in a prophylactic setting. To this end, classical splenic DC (cDC) or DC differentiated *ex vivo* from blood monocytes (MoDC) were pulsed with MHC-I restricted melanoma antigens gp100 and TRP1 and injected s.c. twice, six days apart, into naïve syngeneic mice (illustrated in Fig. 1a). Mice were then challenged with B16F10 tumors, and tumor growth was monitored. Consistent with previous reports^26–28^, significant tumor growth inhibition was observed in vaccinated mice (Fig. 1b). Furthermore, adoptive transfer of splenic CD8^+^ T cells from vaccinated mice protected naïve mice from B16F10 challenge (Fig. 1c), indicating that tumor-reactive CD8^+^ T cells are generated following prophylactic DC vaccination. Next, we tested DC vaccines in an adjuvant setting for their capacity to prevent tumor recurrence. To this end, mice were challenged with B16F10 tumors and allowed to grow until they reached an average size of approximately 100 mm^2^. Tumors were then removed surgically, and mice were left as such, or treated s.c. with cDC or MoDC pulsed with proteins derived from tumors of corresponding mice (illustrated in Fig. 1d). Consistent with clinical reports^17,18^, in the adjuvant setting, both cDC and MoDC protected mice from tumor challenge reoccurrence (Fig. 1e).

**Figure 1:**
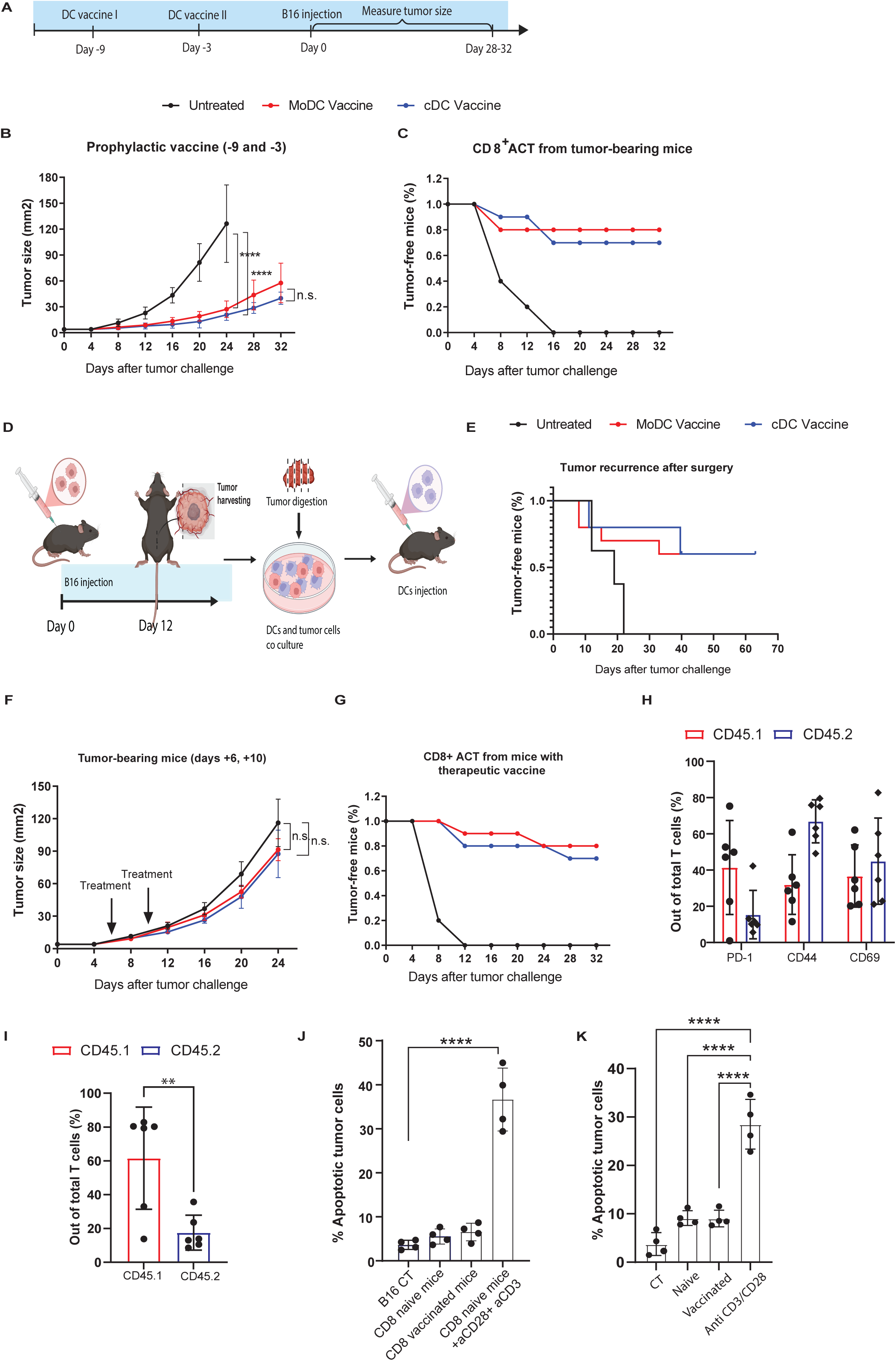
DC vaccines fail to eradicate established tumors despite induction of reactive T cell clones in the draining lymph nodes. (A)Prophylactic vaccine experimental design: Splenic conventional DC (cDCs) and monocyte-derived dendritic cells (MoDCs) cultured from blood were pulsed with gp100 and TRP1 (10 μM each), and injected s.c. twice into naïve syngeneic mice. Mice were then challenged with 2 × 10^5^B16F10 melanoma cells. (B) B16F10 tumor growth in mice that received prophylactic vaccination of cDCs or MoDCs pulsed with melanoma antigens, on days -9 and -3 prior to tumor challenge (n=5). (C) B16F10 tumor-free survival in mice that received 2 × 10⁶ CD8⁺ T cells from either naïve donors or donors vaccinated with prophylactic cDC- or MoDC-based vaccines, following tumor challenge (n=5). (D) Experimental design for B16F10 tumor recurrence: mice underwent surgical removal of B16F10 tumors, followed by a one-week recovery period and then administration of prophylactic cDC-or MoDC-based vaccines to evaluate tumor recurrence. This figure was created using BioRender (https://biorender.com/). (E) B16F10 tumor-free survival in mice that underwent surgical tumor removal followed by prophylactic vaccination with either cDC- or MoDC-based vaccines (n=5). (F) B16F10 tumor growth in mice that received vaccination of cDCs or MoDCs pulsed with melanoma antigens, on days 6 and 10 after to tumor challenge (n=4). (G) B16F10 tumor-free survival in recipient mice that received 2 × 10⁶ CD8⁺ T cells isolated from donor mice previously challenged with B16F10 tumors, with or without vaccination (days +6 and +10) using cDC- or MoDC-based vaccines (n=5). (H)Mean percentages of Caspase 3/7 positive B16F10 melanoma cells after incubation with CD8⁺ T cells isolated from the spleens of naïve C57BL/6 mice or mice vaccinated with activated MoDCbased prophylactic vaccine (n=4). T cell activation (I) pecentge and (J) markers of CD8⁺ T cells from CD45.1⁺ donor mice vaccinated with prophylactic cDC- or MoDC-based vaccines, detected in B16F10 tumors of CD45.2⁺ recipient mice 48 hours after adoptive transfer (n=5). (K) Mean percentages of Caspase 3/7 positive B16F10 melanoma cells after incubation with gp100-specific CD8⁺ T cells isolated from the spleens of naïve mice or mice vaccinated with gp100-pulsed MoDC-based prophylactic vaccine (n=4). Shown is the summary of one experiment out of at least three indecent experiments. Statistical significance was calculated using ANOVA with Tukey’s correction for multiple comparisons (** denotes p<0.001, **** denotes p<0.00001). Error bars represent standard error.

In contrast, however, vaccination of mice-bearing established B16F10 tumors with gp100- and TRP1-pulsed DC did not inhibit tumor growth (Fig. 1f). Nonetheless, adoptive transfer of splenic CD8^+^ T cells from these vaccinated mice protected naïve mice from B16 challenge, indicating that DC vaccines did generate tumor-reactive T cells (Methods, Fig. 1g).

One possibility is that changes in the microenvironment of established tumors limit the infiltration of reactive T cells. However, analysis of adoptively transferred splenic CD8^+^ T cells from vaccinated CD45.1^+^ mice to congenic, tumor-bearing CD45.2^+^ mice, indicated that CD8^+^ T cells do infiltrate into tumors in large quantities and can adapt an active phenotype (Fig. 1hi). Yet, incubation of splenic CD8^+^ T cells from vaccinated mice *in vitro* with B16F10 cells pre-activated with IFNγ (to increase MHC-I levels) and pulsed with the same antigens resulted in only marginal tumor cell killing (Fig. 1j). Since this may be the result of limited tumorreactive clones in wild-type (WT) vaccinated mice, we next used splenic CD8^+^ T cells originating from transgenic mice expressing a gp100-restricted TCR following vaccination with gp100-pulsed cDC. While tumors do not develop in gp100 vaccinated mice (not shown), when we incubated *in vitro* isolated splenic CD8^+^ T cells from these mice, we observed only a marginal B16F10 lysis (Fig. 1k).

### CD8^+^ T cells from tumors and draining lymph nodes bear distinct phenotypes and killing properties

We next set out to understand why CD8^+^ T cells generated by DC vaccines can confer tumorprotecting immunity *in vivo*, but do not kill tumor cells directly *in vitro.* To this end, we adoptively transferred splenic CD8^+^ T cells from gp100 mice, vaccinated with gp100-pulsed DC, to CD45.1 mice bearing B16F10 tumor. After 48 hours, we collected the transferred T cells from the spleen, lymph nodes (pool of axillaries, inguinal, thoracic and mesenteric) and tumors, and assess their killing capacities in vitro. We found that tumor-infiltrating T cells, but not T cells from peripheral sites, induced significant tumor cell lysis upon incubation with tumor cells (Fig. 2a). Additionally, we sorted the antigen-experienced effector CD8^+^ T cells from the DLN and tumors generated spontaneously by B16F10-bearing mice and incubated them *in vitro* with B16F10. Consistently, only effector cells from tumors induced significant tumor cell lysis (Fig. 2b).

**Figure 2:**
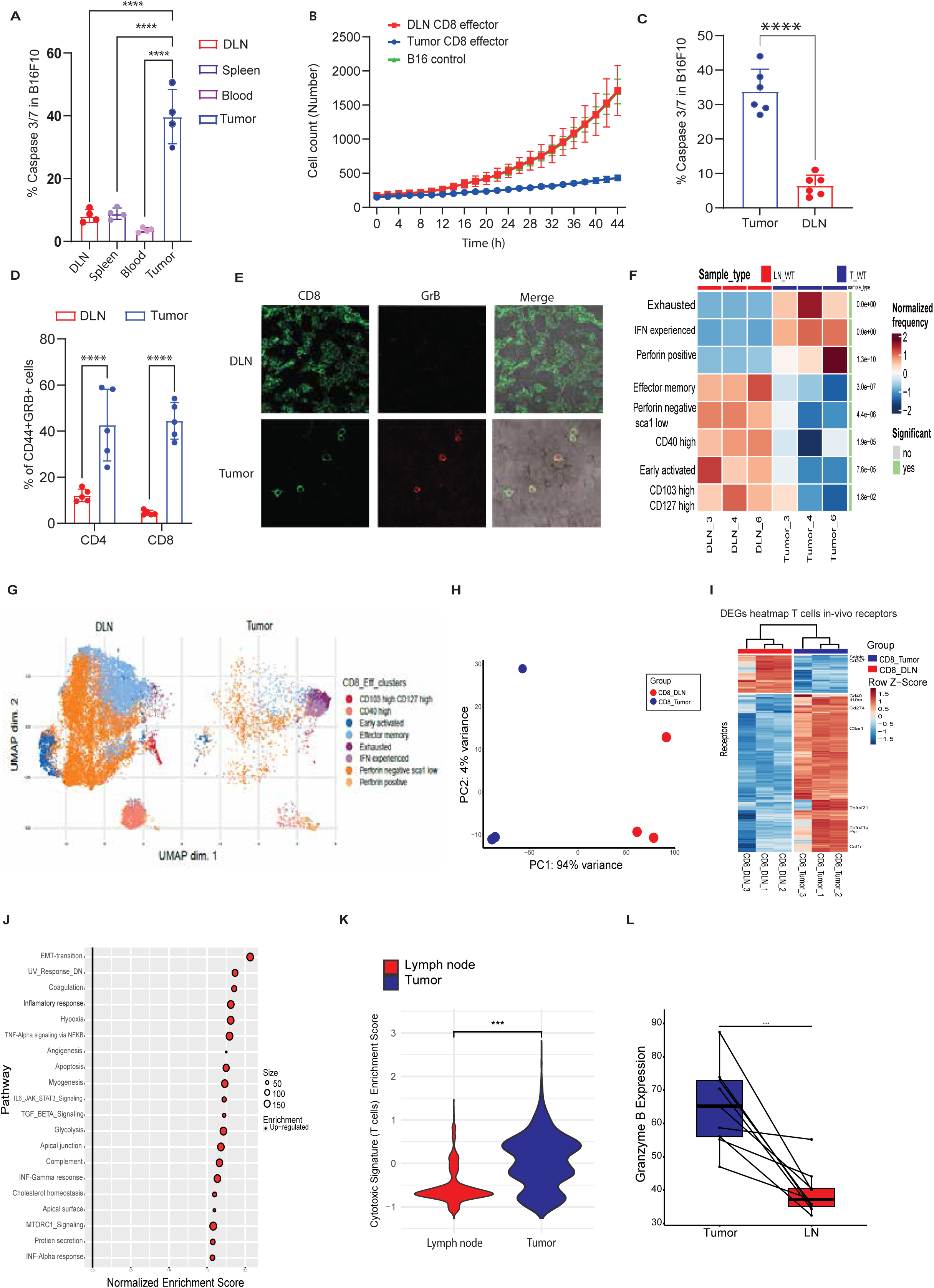
CD8^+^ T cells from tumors and draining lymph nodes bears distinct cytotoxic characteristics and killing properties. (A) Percentages of Annexin V positive tumor cells following over-night incubation with gp100 CD8^+^ T cells (n=10). (B) B16F10 tumor cell counts under IncuCyte following incubation with effector CD8^+^ T cells isolated from LN or tumor of tumor-bearing mice (n=5). (C) Percentages of B16F10 cells positive for Annexin V following overnight incubation with gp100-reactive T cells (n=7). (D) Flow cytometry analyses of GrB expression in gp100-reactive CD8 + T cells in mice bearing B16F10 tumors (n=5). (E) Histological sections of GrB expression in gp100-reactive CD8^+^ T cells in mice bearing B16F10 tumors (n=5). (F) Differential abundance (DA) made by diffcyt of effector CD8^+^ T cells sorted from DLN and tumor of tumor bearing mice. (n=3). (G) UMAP plot of pan CD8^+^ T cells isolated from tumor and DLN of tumor bearing mice (n=3), each point represents an individual cell, colored and annotated by cluster identity. (H) Principal component analysis of gene expression patterns comparing tumor and DLN effector CD8^+^ from tumor bearing C57BL/6 mice (n=3). (I) Hierarchal cluster of significantly changed genes of DLN and tumor CD8+ T cells isolated from B16F10 tumor bearing mice (n=4). (J) Gene set enrichment analysis for all the hallmark gene sets of CD8^+^ T cells from DLN vs TME isolated from B16F10 tumor bearing mice (n=4). (K) Violin plot showing the cytotoxicity signature scores across tumor and LN-derived effector CD8+ T cells (n=9). (L) Granzyme B expression in CD8^+^CD69^+^ T cells from paired tumor and LN samples. Each line connects paired values from individual samples, illustrating differences in expression between tumor and LN tissues (paired t-test, *t* (8) = -5.1, *p* < 0.00093, *n* = 9). Shown is the summary of one experiment out of at least three indecent experiments. Statistical significance was calculated using ANOVA with Tukey’s correction for multiple comparisons (** denotes p<0.001, **** denotes p<0.00001). Error bars represent standard error.

To study whether this may result from differences in abundance of tumor-reactive clones we next isolated CD8^+^ T cells from tumor and DLN of transgenic mice expressing TCRs specific to ovalbumin (OT-I mice) and gp100 that were implanted with B16F10 tumors 10 days prior. Significant tumor cell lysis and granzyme B secretion were measured only when tumorinfiltrating CD8^+^ T cells were incubated with tumor cells (Fig. 2c–2e, Supplemental Fig. 1a).

To shed more light into these differences, we sorted antigen-experienced CD8^+^ T cells from tumor and DLN of day 10 B16F10 melanoma-bearing mice and analyzed their activation phenotype by multiparameter mass cytometry (CyTOF) analysis. In tumors, the majority of CD8^+^ T cells were clustered into exhausted, IFNγ-experienced, or perforin^hi^ T cells. In contrast, the majority of antigen-experienced CD8^+^ T cells in the DLN were mainly effector memory and early activated perforin-negative cells (Fig. 2f–2g and Supplemental Fig. 1b). Additionally, we analyzed the transcriptome of the antigen-experienced CD8^+^ T (SSC_lo_/FCS_lo_/Lin_(CD11b/CD19/B220/NK1.1)neg_/CD3_+_/TCRβ_+_/CD8_+_/CD44_+_/CD62L_neg_) sorted from the DLN and tumor of day 10 tumor-bearing mice by RNAseq. Consistent with our CyTOF data, principal component analysis (PCA) and differentially-expressed gene (DEG) analysis indicated that antigen-experienced CD8^+^ T cells from the DLN and tumors have distinct transcriptional programs (Fig. 2h–2i and supplemental Fig. 1c-1e). Pathway analysis indicated that tumor-infiltrating CD8^+^ T cells exhibit a distinct expression pattern characterized by inflammatory response and high expression of perforin and granzyme B (Fig.2j and Supplemental Fig. 1f).

Next, we determined whether the findings observed in mice regarding the states of CD8^+^T cells and DC subsets in the TME and lymph nodes (LNs) were also reflected in humans. To achieve this, publicly available human scRNA-seq data from patients with HNSCC were reanalyzed^29^, focusing on the expression profiles and functional states of CD8^+^ T cells in matched tumors and LNs. The analysis revealed 7 distinct clusters with different proportions of cells originating from the tumors or the LNs (Supplemental Fig. 2a-2b). Clusters were manually annotated based on their differentially expressed (DE) genes. To align with the mouse experiments, only the effector CD8^+^ T cell clusters were analyzed, which are characterized by high expression of activation and cytotoxicity-related genes, including *PRF1*, *CD44*, *CD69*, *HSPA1A*, *CXCL13*, *TNFRSF18*, and *TYROBP*.

Comparing differentially expressed genes between tumor-derived and LN-derived effector CD8^+^ T cells revealed significant upregulation of cytotoxic genes, such as *GZMB* and *PRF1*, in tumor-derived cells, indicating increased cytotoxic potential. Additionally, genes associated with T cell exhaustion were also upregulated, highlighting the dual nature of activation and exhaustion in tumor-infiltrating CD8^+^ T cells (Supplemental Fig. 2c-2e). To validate these observations, cytotoxic gene signature enrichment scores were calculated. Indeed, tumorderived effector CD8^+^ T cells exhibit significantly higher cytotoxic activity compared to LNderived effector CD8^+^ T cells (Fig. 2k).

Moreover, CyTOF data from the same study were reanalyzed. The gating strategy mirrored that used in the mouse experiments, focusing on effector CD8^+^ T cells. Expression values for granzyme B were then assessed in CD8^+^/PD1^+^ and CD8^+^/CD69^+^ T cells from both tumor and LN tissues. The results revealed a higher proportion of activated CD8^+^ T cells in the tumor compared to the DLN. Additionally, Granzyme B expression was elevated in tumor-infiltrating CD8^+^ T cells relative to those from the LN. Indeed, the analysis showed significant difference in the proportions of activated CD8^+^ T cells between tumor and LN samples, further supporting the heightened activation and cytotoxicity of tumor-infiltrating CD8^+^ T cells (Fig. 2l).

### Distinct dendritic cell (DC) populations dominate the tumor microenvironment and lymph nodes

Next, we assessed the process by which antigen-experienced CD8+ T cells in the DLN differentiate into cells with a cytotoxic phenotype in tumors. Findings from our and others’ studies suggested that cues from tumor-infiltrating DC (TIDC) may play a role in that process^10,30–32^. To address that possibility, we sorted antigen-experienced CD8^+^ T cells (SSC_lo_/FCS_lo_/Lin_(CD11b/CD19/B220/NK1.1)neg_/CD3_+_/TCRβ_+_/CD8_+_/CD44_+_/CD62L_neg_) from the DLN of day 10 B16F10 tumor-bearing mice and incubated them *in vitro* with B16F10 and with either pan DC sorted from the DLN or with TIDC. Indeed, significantly higher killing rates were observed in CD8^+^ T cells upon addition of TIDC, but not DLN DC (Fig. 3a). Next, we aimed to characterize the DC subsets that occupy the tumor and DLN. To this end, B16F10 melanoma cells were injected into C57BL/6 mice and Lin^(CD3/NK1.1/CD19)neg^/CD11c^+^/MHCII^hi^ cell populations were analyzed by CyTOF. In tumors, the vast majority of DCs were from the myeloid lineage, including dermal DC subsets and cDC2 expressing CD11b. In contrast, DLN were mainly populated by classical DC subsets and minor populations of myeloid DC (Fig. 3b–3c and supplemental Fig 3a-3b). Consistently, flow cytometric analysis of day 10 B16F10 tumor-bearing mice further corroborated that MoDCs specifically occupy the tumor sites, whereas cDCs were more prevalent in lymphoid organs across mice (Fig. 3d). To assess the human relevance of these patterns, publicly available human scRNA-seq data from patients with HNSCC were utilized. Clusters with high CD11c expression were further analyzed to compute the proportions of cDCs and MoDCs in both tumor- and LN-derived cells. Enrichment scores of MoDC and cDC gene signatures in the tumor and LN were calculated. The analysis revealed that MoDCs were significantly more enriched in the TME compared to the LN, suggesting their potential role in the secondary activation of CD8^+^ T cells within the tumor. In contrast, cDCs were more enriched in the LN than in the TME, indicating their role in the initial activation of CD8^+^ T cells and the regulation of immune responses in lymphatic tissues (Fig. 3e–3f).

**Figure 3:**
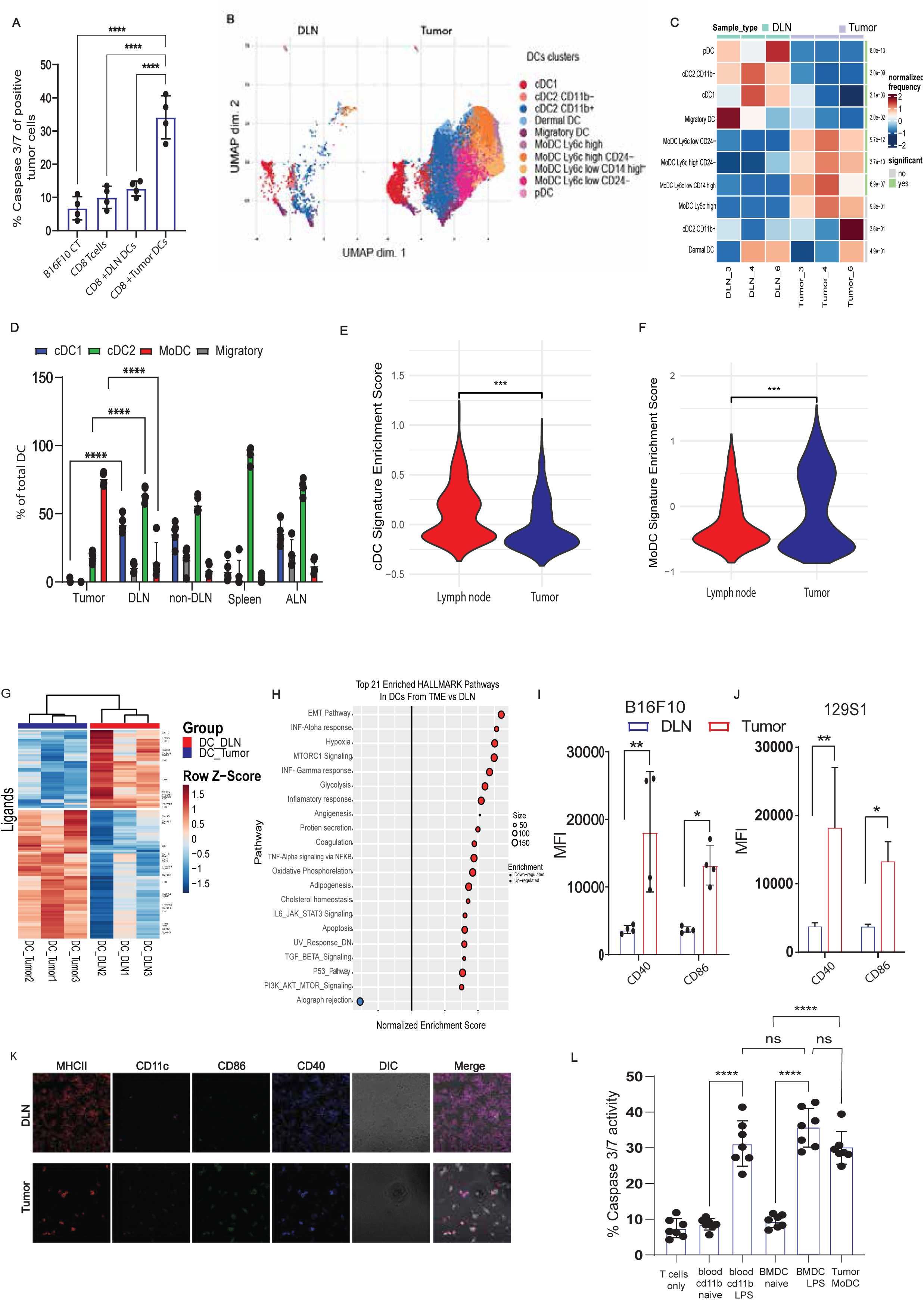
Tumor-infiltrating DC and lymph nodes-resident DC are occupied by different DC populations. (A) Percentages of B16F10 cells positive for caspase 3/7 following overnight incubation with DLN CD8+ T cells plus DLN/Tumor-DC (n=4). (B) UMAP plot of pan CD11C^High^ MHCII^high^ cells isolated from tumor and DLN of tumor bearing mice (n=3), each point represents an individual cell, colored and annotated by cluster identity. (C) Differential abundance (DA) made by deficit of CD11C^High^ MHCII^high^ cell populations isolated from DLN and tumor of tumor bearing mice(n=3). (D) Mean percentage of dendritic cell (DC) subsets in various organs of B16F10 tumor-bearing mice, analyzed on day 10 after tumor challenge (n=3). (E) Violin plot showing the cDC signature scores across tumor-derived and LN-derived cells. (F)Violin plot showing the MoDC signature scores across tumor-derived and LN-derived cells. (G)Hierarchal cluster of significantly changed genes in tumor vs DLN DCs isolated from B16F10 tumor bearing mice (n=4). (H) Gene set enrichment analysis for all the hallmark gene sets of DCs cells from DLN vs TME isolated from B16F10 tumor bearing mice (n=4). (I-J) Mean fluorescence intensity (MFI) of activation markers CD40/CD86 of DCs isolated from the DLN and tumor of (I) C57BL/6 tumor bearing mice (n=4), (J) 129S1 mice (n=3). (K) Histological sections of CD40/CD86 expressing DCs in mice bearing B16F10 tumors (n=4). (L) Percentages of B16F10 cells positive for caspase 3/7 following overnight incubation with CD8^+^ T cells plus naïve and LPS actiavted MoDCs from different organs vs TME (n=4). Shown is the summary of one experiment out of at least three indecent experiments. Statistical significance was calculated using ANOVA with Tukey’s correction for multiple comparisons (** denotes p<0.001, **** denotes p<0.00001). Error bars represent standard error.

To assess potential differences in gene expression, we next sorted the FCS ^lo^/SSC^lo^/Lin^(CD3/NK1.1/CD19)neg^/CD11c^+^/MHCII^hi^ cells from the tumor and DLN of day 10 B16F10-bearing mice and analyzed their transcriptome by bulk RNAseq. PCA showed clear differences between tumor-derived and DLN-derived DCs (Supplemental Fig.3b). These transcriptional differences were further emphasized by focusing on expressed ligands, which in part reflect the immune regulatory potential of the two DC subsets (Fig. 3g). Notably, TIDC manifested a distinct inflammatory response, characterized by production of inflammatory cytokines (i.e. IL-6, IFN, TNF), chemokines (i.e. CXCL10, CXCL1) and costimulatory molecules (Fig. 3g–3h and supplemental Fig. 3c-3e). Flow cytometric analyses and histological sections of tumoral and DLN DCs indicated that the former exhibit significantly higher levels of co-stimulatory molecules in both syngeneic mice (B6) and in mice mounting tumoreradicating immunity (129S1)(Fig. 3i–3k). Analysis of multiple tumor mouse models indicated that the phenotype of MoDC was significantly more active, across spontaneous (MMTV-PyMT and Apc^min^ mice) and transplanted tumor models (4T, MC38), suggesting that this activation pattern represents a broader and generalizable phenomenon (Supplemental Fig. 3f-3k).

Next, we tested whether the capacity of TIDC to activate antigen-experienced CD8^+^ T cells is a result of activation patterns unique to tumors or could be replicated outside the tumor context. To this end, we isolated monocytes from the blood and BM of naïve mice and matured them *in vitro* to DC. We then added them as such, or following overnight LPS activation, to a co-culture of CD8^+^ T cells isolated from the DLN of tumor-bearing mice and B16F10 tumor cells. The effect of naïve monocyte-derived DC from both BM- and blood on the tumor-killing ability of CD8^+^ T cells was negligible. In contrast, LPS-activated MoDC significantly increased tumor cell killing by CD8^+^ T cells, comparable to the effect observed by TIDC (Fig. 3l).

### CD8^+^ T cell cytotoxicity requires successive cues from classical and myeloid DC

In order to identify the cues required for CD8^+^ T cells to elicit cytotoxicity, we next established an *in vitro* experimental system. Initially, naïve T cell (CD62L^+^/CD44^neg^) clones recognizing gp100_25-33,_ or OVA_257-264_ peptides were incubated overnight with B16F10 or MC38 tumor cells pulsed with the corresponding peptides (Fig. 4a). As indicated by caspase 3/7 activity in tumor cells, we found that incubation of T-cell clones does not result in significant tumor cytotoxicity over baseline (Fig. 4b). To test if priming by LN cDC could endow CD8^+^ cytotoxic activities, we next incubated the isolated gp100_25-33,_ or OVA_257-264_ -reactive CD8^+^ T cells with antigenpulsed cDC for three days. CD8^+^ T cells were then isolated from the culture and incubated overnight with the corresponding antigen-pulsed tumor cells (Fig. 4a). Similar to our results with freshly isolated gp100_25-33,_ or OVA_257-264_ reactive CD8^+^ T cells, we did not observe significant tumor cell killing over baseline (Fig. 4b). In sharp contrast, when we added LPSactivated MoDC (1:1 ratio) to the co-culture of CD8^+^ T cells pre-incubated with cDC, we observed a potent tumor cytotoxicity (Fig. 4c–4d).

**Figure 4:**
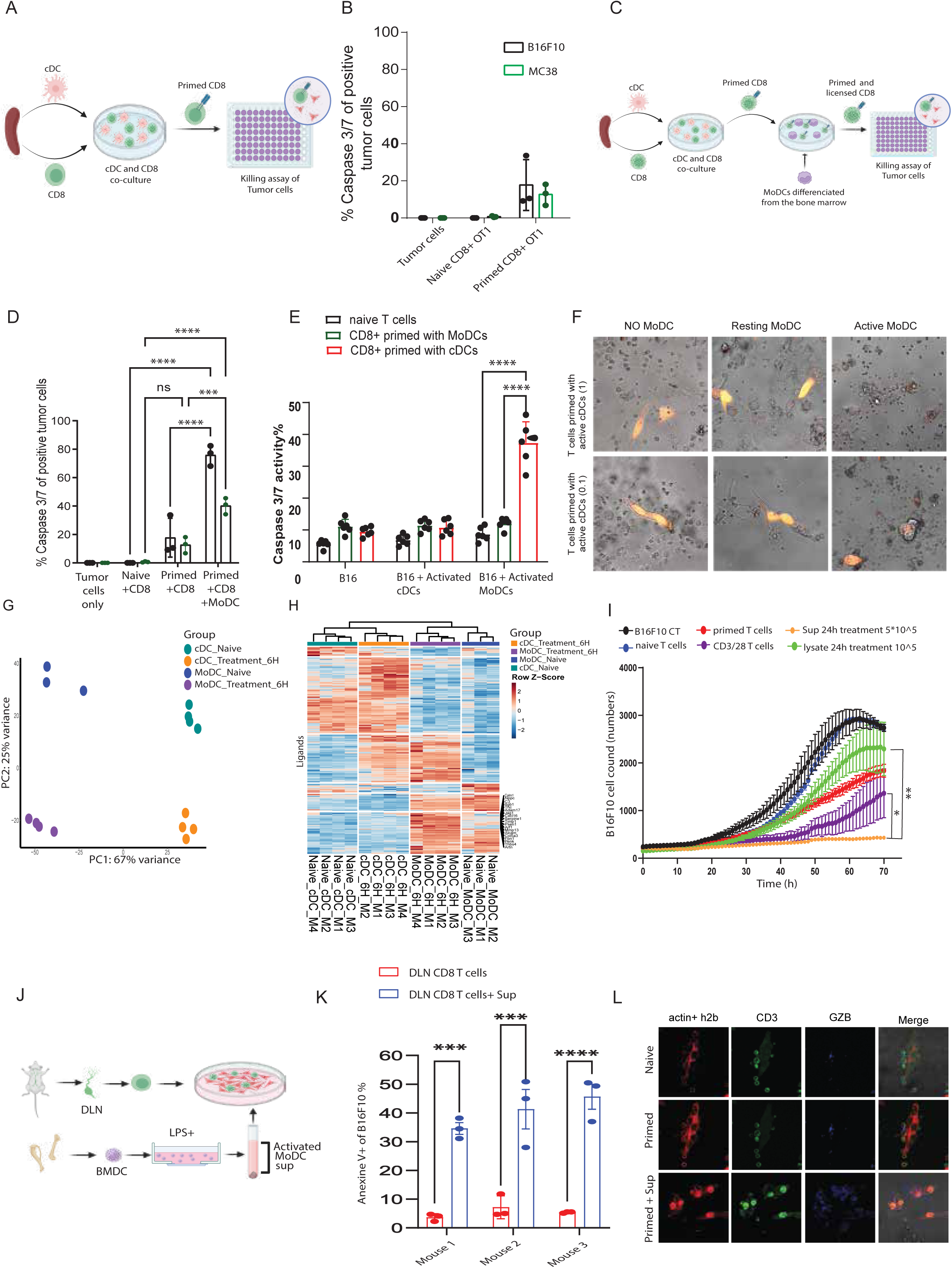
CD8^+^ T cells cytotoxicity requires subsequent cues from classical and myeloid DC. (A) Illustration of experimental outline (https://biorender.com/). (B) Percentages of B16F10/MC38 OVA loaded positive cells for caspase 3/7 following overnight incubation with antigen experienced CD8+ T cells (n=3. (C) Percentages of B16F10/MC38 OVA loaded positive cells for caspase 3/7 following overnight incubation with antigen experienced CD8+ T cells and LPS-activated MoDC (n=3). (D) Illustration of experimental outline (https://biorender.com/). (E) Percentage of B16F10 tumor cells positive for caspase 3/7 following incubation with gp100 CD8 T cells primed with cDCs or MoDCs with the addition of LPS-activated cDC or MoDC (n=6). (F) Representative confocal images of B16F10-OVA cells incubated overnight with OTI- CD8 T cells and MoDCs. (G) Principal component analysis of gene expression patterns comparing cDC and MoDC following activation with LPS for 6 hours (n=3). (H) Hierarchal cluster of significantly changed genes in cDC and MoDC following activation with LPS for 6 hours (n=4). (I) IncuCyte counts of live B16F10 cells overtime following incubation with CD8^+^ T cells plus sup/lysate of activated MoDCs. (J) Illustration of experimental outline (https://biorender.com/). (K) Percentages of B16F10 cells positive for Annexin V following overnight incubation with DLN CD8^+^ T cells plus LPS activated MoDCs sup (3biological repeats and 3 technical repeats each). (L) Representative confocal microscopy image of GrB expression in DLN CD8^+^T cells incubated overnight with B16F10 cells w/o activated MoDCs sup. Shown is the summary of one experiment out of at least three indecent experiments. Statistical significance was calculated using ANOVA with Tukey correction for multiple comparisons. *, P < 0.05; **, P < 0.01; ****, P < 0.0001.

We therefore hypothesized that cDC and MoDC may provide complementary, sequential signals to T cells that elicit their cytotoxic capacity. To test this hypothesis, we isolated naïve splenic CD8^+^ T cells from gp100_25-33_ or OVA_257-264_ reactive TCR mice and incubated them for 3 days with corresponding antigen-pulsed cDC or MoDC. CD8^+^ T cells were then isolated and added to a culture of B16F10 tumor cells with either LPS-activated cDC or MoDC (illustrated in Fig. 4c). We found that only CD8^+^ T cells pre-activated with cDC and later incubated with activated MoDC elicit significant cytotoxicity (Fig. 4e–4f). Indeed, analysis of the transcriptional profiles in cDC and MoDC indicated that the two subsets express a distinct set of genes at baseline level (Fig. 4g–4h) and following activation with LPS, MoDC exhibited a unique and much stronger inflammatory response (Fig. 4h and supplemental Fig. 4b). Next, we set out to identify the signal(s) by which MoDC induces CD8^+^ T cells’ cytotoxicity, potentially a surface receptor or secreted mediator. To this end, we activated overnight MoDC with LPS and added the supernatants, or cell lysates, to a co-culture of cDC-activated gp100_2533_ reactive CD8^+^ T cells and B16F10 cells, after which we measured the killing rates overtime under IncuCyte. Our results indicated that supernatant, but not cell lysates, induced tumor cell lysis (Fig. 4i). To corroborate these findings, we isolated gp100_25-33_ reactive CD8^+^ T cells from the DLN of transgenic mice bearing B16F10 tumors and incubated them *in vitro* with B16F10 tumor cells, as such or with supernatants of activated MoDC (Fig. 4j). In agreement with our previous data, addition of MoDC supernatants induced tumor cell lysis and granzyme B secretion by CD8^+^ T cells (Fig. 4k–4l).

### MoDC activate primed CD8^+^ T cells through TNFR signalling to elicit tumor cytotoxicity

We next set out to identify the potential ligand-receptor interactions that license the cytotoxic activity of CD8^+^ T cells. First, we aimed to identify genes predominantly expressed in T cells following priming with cDC and activation with MoDC. To this end, we incubated naïve gp100_25-33_ TCR CD8^+^ T cells, with cDC and MoDC pulsed with gp100 peptide for three days and analyzed their transcriptome. Additionally, we isolated CD8^+^ T cells after 3 days of priming with cDC and activated them for 6 hours with supernatants of activated MoDC and cDC and analyzed changes in their gene expression profile (illustrated in Fig. 5a). Priming of CD8^+^ T cells with cDC followed by activation with supernatants from MoDC resulted in a distinct expression pattern characterized by increased chemokines and inflammatory cytokine signalling (Fig. 5b–5c and supplemental Fig. 5a).

**Figure 5:**
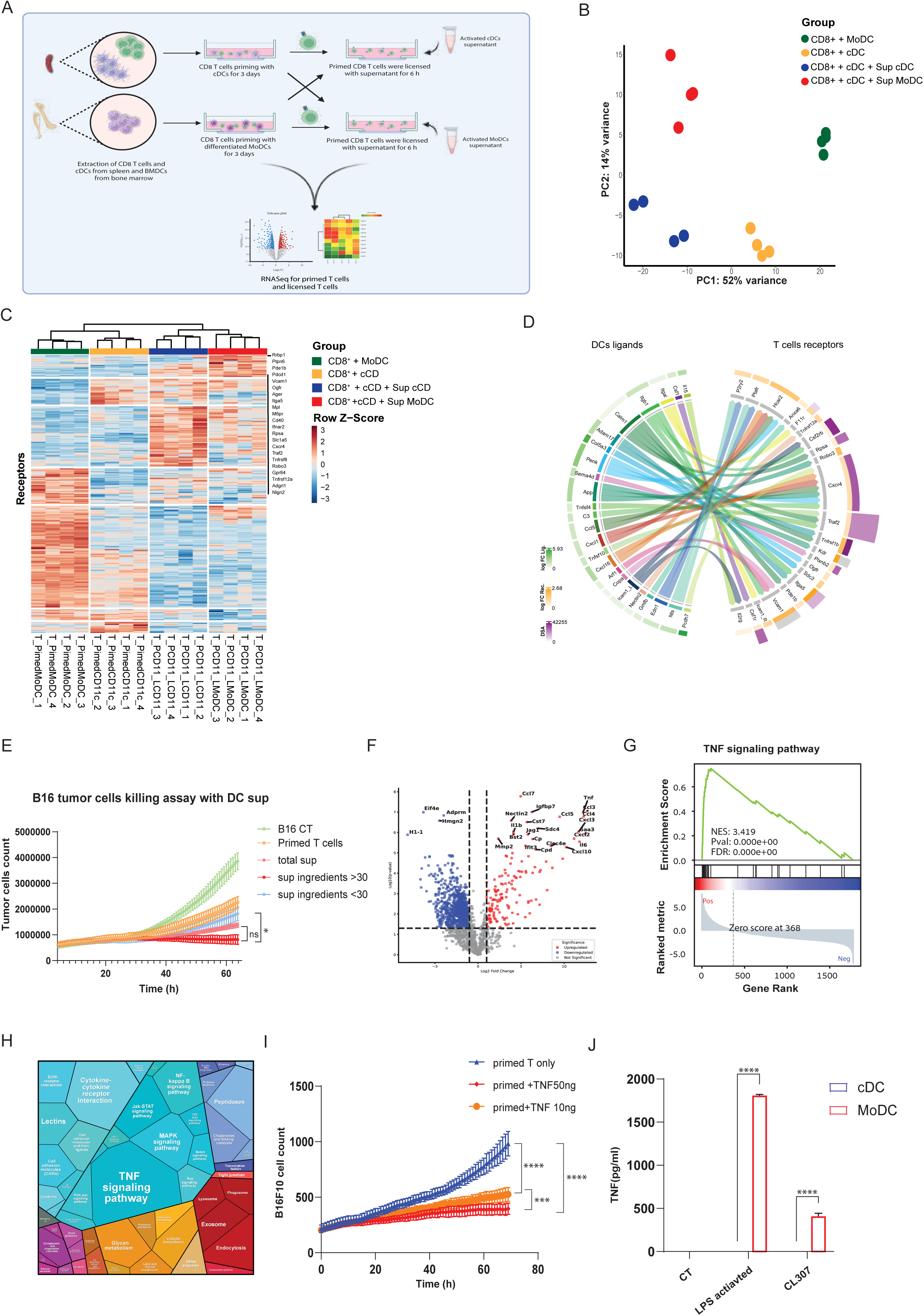
MoDC activate primed CD8^+^ T cells through TNF signaling to elicit tumor cytotoxicity. (A) Illustration of experimental outline (https://biorender.com/). (B) Principal component analysis of gene expression patterns comparing different combinations of primed (P) and licensed (L) CD8^+^ T cells by cDCs or MoDCs (n= 4). (C) Hierarchal cluster of significantly changed genes in primed (P) and licensed (L) CD8^+^ T cells (T) by cDCs or MoDCs (n= 4). (D) Circus plots showing connecting unique MoDC ligands with receptors on antigen-experienced T cells (n=4). (E)B16F10 tumor cell counts in IncuCyte following incubation with effector CD8 + T cells and sup fraction of LPS activated MoDC. (F) Volcano plot showing differential expression of proteins between 6-hour LPS-activated MoDCs and Naive MoDCs (>30 k Da fraction) (n=3). (G) GSEA enrichment plot for the “TNF Signalling Pathway,” showing the distribution of leading-edge genes along the ranked gene list. The plot highlights the correlation between gene rank and pathway enrichment, with the running enrichment score (ES) shown on top. The vertical line marks the position of the leading edge (n=3). (H) Proteomap (level 3) showing enriched signalling pathways from significantly and non-significantly upregulated proteins in activated MoDCs (>30 kDa fraction) (n=3). (I) IncuCyte live-cell counts of B16F10 melanoma cells over time following incubation with OVA-primed CD8^+^ T cells with or without the addition of TNF. Shown is the summary of one experiment out of at least three indecent experiments. (J) ELISA analysis of TNF-alpha cytokine productions by naïve, LPS or CL307 actiavted MoDCs vs classical DCs, differentiated from bone marrow and the spleen of C57BL/6 mice, respectively (n=3). Statistical significance was calculated using ANOVA with Tukey correction for multiple comparisons. *, P < 0.05; **, P < 0.01; ****, P < 0.0001.

To dissect the ligand-receptor interactions that license the cytotoxic activity of CD8^+^ T cells, we focused on receptors specifically upregulated in CD8^+^ T cells primed with cDCs and licensed with MoDCs compared to CD8^+^ T cells primed with cDCs alone. These receptors were further associated with ligands that were uniquely upregulated in activated MoDCs (illustrated in supplemental Fig. 5b,). To quantify the potential influence of these interactions, we calculated the intracellular downstream activation (DSA) values for each receptor using InterFlow algorithm^33^. DSA values, which reflect the impact of each receptor’s activation on downstream signalling pathways in CD8^+^ T cells. Higher DSA values indicate a stronger influence of the ligand-receptor interaction on CD8^+^ T cell functionality. Notable receptors with high DSA values include *Traf2*, *Tnfrsf1b*, *Cxcr4*, and *Csf1r* (Fig. 5d).

To narrow the list of potential ligands, we next separated the MoDC supernatant (sup) on size exclusion columns and tested the different fractions for their capacity to activate CD8^+^ T cells against B16F10 tumor cells. Incubation of CD8^+^ T cells with the >30 kDa supernatant fraction exhibited significantly higher rates of tumor cell killing compared to the <30 kDa supernatant fraction (Fig. 5e).

Next, proteins from the >30 kDa fraction of activated and naïve MoDCs were purified and subjected to proteomic analysis. A volcano plot comparing activated and naïve MoDCs revealed several significantly upregulated proteins (adjusted p-value < 0.05, |log2(fold change)| > 1), including *Tnf* (Fig. 5f). Gene Set Enrichment Analysis (GSEA) enrichment for the TNF signalling pathway demonstrated a strong positive enrichment score, peaking early in the ranked gene list, indicating that the genes contributing to this pathway are predominantly located at the top of the ranked list (Fig. 5g). The upregulated proteins in activated MoDCs were analyzed using the Proteomaps tool to identify enriched pathways and biological processes. The resulting map highlights the TNF signalling pathway as the most enriched, alongside significant contributions from the cytokine-cytokine receptor interaction and MAPK signalling pathways (Fig. 5h).

To validate the impact of TNF on the cytotoxic activity of CD8^+^ T cells, we isolated splenocytes that were extracted from the spleens of gp100 mice and incubated with gp100 peptide -pulsed cDC for 3 days. Next, T cells were isolated and subsequently incubated with B16F10 cells for 72 hours. Primed CD8^+^ T cells were treated with varying concentrations of murine TNF ligands. TNF-treated, primed CD8^+^ T cells exhibited significantly greater tumor cell killing compared to untreated, primed CD8^+^ T cells (Fig. 5i and Supplemental Movies1-5). Consistently, analysis of the supernatant of LPS-activated cDC isolated from the lymph nodes secrete only negatable levels of TNFa. In sharp contrast, MoDC cultured from blood monocytes secrete high levels of TNFa upon overnight stimulation with LPS (Fig. 5j).

### DC rapidly lose their capacity to license the cytotoxic activity of primed CD8^+^ T cells

Consistent with the reduction of MoDC observed overtime in tumors (Supplemental Fig. 6a), we also found a significant reduction in the levels of TNFα (Fig. 6a). Hence, we speculated that injection of activated MoDC into tumors could restore the levels of TNFα required to support cytotoxicity. To this end, we isolated monocytes from CD45.1^+^ mice and matured them *ex vivo* to obtain MoDC. Next, MoDC were activated overnight with LPS, pulsed with MHC-I restricted gp100 peptide and injected s.c into CD45.2^+^ congenic mice bearing palpable B16F10 tumors (Illustrated in supplemental Fig. 6b). After 24 and 48 hours, mice were sacrificed, and the percentages of CD45.1 MoDC were analyzed by flow cytometry. CD45.1^+^MoDC were found in the inguinal draining lymph nodes as well as the spleen. About 8 percent of the total tumor DC were CD45.1^+^/CD45.2^neg^, demonstrating their capacity to infiltrate tumors (Fig. 6b). Nonetheless, analysis of their activation phenotype in both the tumor bed and the lymph nodes revealed that they had almost completely lost their expression of TNFα and costimulatory molecules within 24 h following injection (Fig. 6c–6d and Supplemental Fig. 6a-6b).

**Figure 6:**
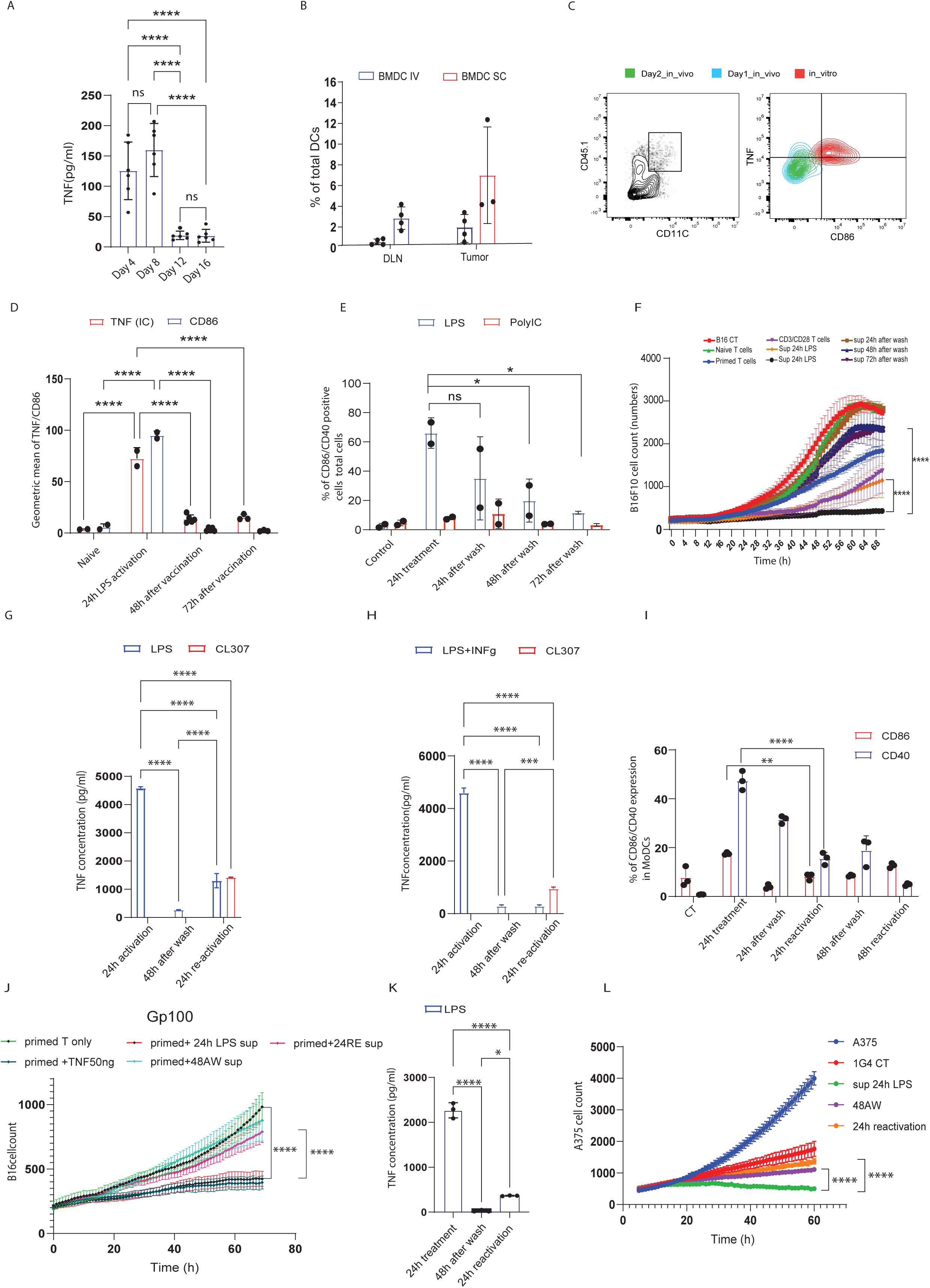
DC rapidly lose their capacity to license the cytotoxic activity of primed CD8^+^ T cells. (A) ELISA analysis of TNF-alpha cytokine productions by cells isolated from the tumor of tumor bearing mice on different days (n=6). (B) Mean percentage of CD45.1 MoDCs detected in B16F10 tumors 48 hours after vaccination. MoDCs were activated overnight with LPS, pulsed with gp100 peptide, and injected either s.c. or i.v. into CD45.2⁺ mice 10 days after tumor challenge (n=3 or 4). (C) FACS analysis of CD45.1 DCs expressing TNF in vitro and CD45.1 DCs isolated from the tumor of B16F10 tumor-bearing C75BL/6 mice (day 10/11; *n* = 5). (D) Mean percentage of CD86 and TNF expression on CD45.1⁺ MoDCs detected in B16F10 tumors at 24- and 72-hours post-vaccination. MoDCs were activated overnight with LPS, pulsed with gp100 peptide, and injected s.c. into CD45.2 mice 10 days after B16F10 tumor challenge (n=2,3,4 or 5). (E) Expression of CD86/CD40 activation markers on MoDCs differentiated from the bone barrow of C57BL/6 mice, after LPS or PolyIC activation and wash. (n=3). (F) IncuCyte live-cell counts of B16F10 melanoma cells over time following incubation with primed CD8+ T cells with or without LPS-activated or washed MoDCs sup (n=3). (G) ELISA analysis of TNF-alpha cytokine productions by mouse MoDCs differentiated from bone marrow of gp100 mice, and underwent LPS treatment, wash and LPS/CL307 reactivation (n=3).. (H) ELISA analysis of TNF-alpha cytokine productions by mouse MoDCs differentiated from bone marrow of gp100 mice, and underwent LPS+INFγ treatment, wash and LPS/CL307 reactivation. (I) Expression of CD86/CD40 activation markers on MoDCs differentiated from the bone barrow of C57BL/6 mice, after LPS/PolyIC activation, wash and reactivation (n=3). (J) IncuCyte live-cell counts of H2b-tdt gp100 loaded B16 melanoma cells over time following incubation with naïve or primed CD8 cells with or without LPS-activated or washed MoDCs sup (n=3). (K) ELISA analysis of TNF-alpha cytokine productions by MoDCs differentiated from blood monocytes of healthy donors, and underwent LPS treatment, wash and reactivation. (L) IncuCyte live-cell counts of A375 human melanoma cells over time following incubation with 1G4 TCR infected CD8 T cells with or without LPS-activated or washed human MoDCs sup. Shown is the summary of one experiment out of at least three indecent experiments. Statistical significance was calculated using Two-way ANOVA with Tukey correction for multiple comparisons. *, P < 0.05; **, P < 0.01; ****, P < 0.0001.

To assess if this phenomenon resulted from the suppressive tumor microenvironment, we also tested their activation phenotype over time *in vitro*. Hence, MoDC were activated overnight with LPS, then washed and incubated with complete media, and analyzed for their activation phenotype overtime. A significant reduction in MoDc expression of costimulatory molecules occurred over time, reaching baseline levels approximately 48 h after wash (Fig. 6e). Consistently, the capacity of their supernatants to support *in vitro* killing of B16F10 by gp100reactive CD8^+^ T cells was completely abrogated 48 hours after LPS was washed out (Fig 6f). Analysis of TNFα secretion at these timepoints demonstrated that their levels in the supernatants dropped to almost zero after LPS wash out and correlated (Fig. 6g). Interestingly, attempts to reactivate MoDC a second time with LPS or with a TLR7/8 ligand (i.e. CL307) did not induce expression of co stimulatory molecules or TNFα secretion (Fig. 6g–6i).

Consistently, the supernatants of reactivated MoDC failed to support the cytotoxic activity of primed gp100-restricted CD8+ T cells (Fig. 6j).

Lastly, we tested the relevance of these findings to human cells. To this end, blood monocytes were isolated from healthy donors and matured in vitro to obtain MoDC and activated with LPS. As expected, large amounts of TNFα were detected in the supernatants (Fig. 6k) and declined almost to baseline levels within 24 hours after LPS was washed out. Most importantly, reactivation of these MoDC resulted in a negligible amount of secreted TNFα (Fig. 6k). To test its effect on the cytotoxic activities of CD8^+^ T cells, we transduced blood CD8^+^ T cells with the 1G4 TCR bearing physiological micromolar affinity to HLAA2:NYESO1 complex and incubated them at a ratio of 1:5 (E:T) with A375 target cells. Although baseline tumor cytotoxicity was observed in this culture, it was significantly potentiated upon addition of supernatant from LPS-activated MoDC. Consistent with the levels of TNFa in these supernatants, addition of supernatants from activated MoDC 48 hours after LPS wash, or of MoDC reactivated with LPS, was almost inert and had no effect on T cells’ cytotoxicity (Fig. 6l and Supplemental movies 6-10).

## Discussion

Over half a century, the trajectory of T cells from naïve to cytotoxic effector cells, and the regulatory signals and cells that govern this process, have been in the center of immunology research. Our current understanding is that antigens must be presented by DC in the context of co-stimulatory molecules and secretion of inflammatory cytokines, and predominantly IL12^2,34^. This context can be induced directly through activation of pattern-recognition receptors^35,36^, or following CD40-CD40L interactions and/or secretion of IFNγ from primed CD4^+^ T cells^37,38,39^. Subsequent studies have demonstrated that cDC1 is the main DC subset that prime both CD4^+^ and CD8^+^T cells in the lymph node^1,40,41^. Indeed, identification of genes predominantly expressed in different DC subsets and the use of corresponding knock-out mouse models have highlighted the critical role of cDC1, which are the main DC capable of cross-presentation, inducing CD8 T cell responses to extracellular antigens^1^.

Along these lines, the discovery of cDC subsets in tumors and their correlation with favourable outcomes has been mainly attributed to their capacity to present tumor antigens in the DLN ^8,42,43^, or in tertiary lymphoid structures^44^. Therefore, most studies have focused and emphasized the migration pattern of cDC and their capacity to stimulate CD8^+^ T cells. Several important studies emphasize the importance of cDC in tumor, and these suggest that intratumoral cDC may act through secretion of CXCL9/CXCL10 to promote T cell infiltration ^45,46^. Other publications suggested that IL-12 can be provided intratumorally by infiltrating cDC to increase T activity_31,47._

However, this model of T cell activation does not provide an explanation for the limited efficacy of DC vaccines. Numerous clinical trials have attempted to employ the presentation of tumor antigens by DC in order to elicit T-cell immunity^12,13^. On one hand, seminal studies have shown that vaccination with neoantigens loaded on activated DC, or injected with TLR3 agonist, elicit strong T-cell immunity and prevent tumor recurrence in patients with resected melanoma ^16–18^. Similarly, mRNA transfected DC showed promising results in the adjuvant setting^48,49^. In contrast, however, attempts to employ DC to treat established tumors, rather than as prophylactic tumor vaccines, have achieved only marginal success in clinical trials^12,19^. While the presence of a suppressive microenvironment in established tumors is likely to inhibit tumor-reactive T cells, clinal data indicate that tumor-reactive T cells do infiltrate into tumors following vaccination ^23,24^.

In addition, several key questions may be raised; for instance, a number of seminal studies demonstrated that TCR-MHC is more promiscuous than earlier thought, allowing each T cell to recognize several to hundreds of different peptides^50,51^. Hence, why does licensing T cells in the lymph node rarely result in off-site cytotoxicity? Why is improved immunity obtained in mice injected intratumorally with immune modulators, but not systemically? Our data suggests that T cell cytotoxicity is elicited through two activation modules that are distinct from one another by the nature of signals and the type of DC subsets that provide them.

Consistent with our previous findings^30^, we suggest here that the main role of tumor-infiltrating MoDC is not to present antigens, but rather to provide a licensing signal to CD8^+^ T cells primed in the DLN by cDC. The importance of intratumoral MoDC for the induction of CD8^+^ T cell immunity was demonstrated in the pioneering work of Kohn and colleagues^52,53^ and was further supported by several important studies. Prokhnevska et al. have suggested tumor-reactive CD8^+^ T cells are activated in the DLN but convert to effector cells only in the tumor as a result of subsequent costiumlation cues at from the microenvironment. Yet, the nature of these cues and the cell subsets providing them remained unclear. Garris and colleagues suggested a similar two-step model of CD8^+^ T cell activation and suggested it is mediated by IL-12 and IFNγ secreted by tumor-infiltrating DC ^31^. Here, we demonstrate that TNFα secreted from MoDC is the main factor eliciting cytotoxicity from primed T cells infiltrating tumors. Sabrina and colleagues were the first to identify a myeloid subset of DC in the spleens of Listeria monocytogenes-infected mice, which produce TNFα and iNOS, and coined the term Tip DC^54^. Tip DCs were also found to be a major source of TNFα in other infectious diseases, including fungi^55^, parasites^56^, and autoimmunity onsets and especially psoriasis^57^. Marigo and colleagues were the first to demonstrate the necessity of Tip DC for the success of adoptive CD8^+^ T cell therapies against tumors and suggested CD40-CD40 ligand interactions as the main mechanism of activation^58^. Yet, the direct effect of TNFα within the context of CD8^+^ T cells remained unclear in these studies.

TNFα was discovered by Dr. Lloyd J. Old and colleagues due to its ability to induce tumor necrosis in tumor-bearing mice^59^. Subsequent studies have supported the crucial role of TNF in mediating T cell tumor immunity, promoting survival, and antigen induced differentiation and activation of T cells. In support of that notion, TNFR2 in tumor infiltrating lymphocytes from patients with triple-negative breast cancer was found to be associated with improved survival^60^. Consistently, agonistic monoclonal antibodies against TNFR2 were shown to significantly improve CD8^+^ T cell immunity in humanized mouse models^61^. Yet, the failure of TNFα in the clinic^62,63^, as well as the improved tumor immunity observed in mice lacking TNFα or both TNF receptors^64,65^, have questioned its favourable role in cancer. Along these lines, TNFα was shown to mediate a range of pro-tumor activities including angiogenesis, elevation of checkpoint blockade receptors, and proliferation of cancer fibroblasts and epithelial cells^66^. Our results may suggest that the paradoxical roles reported for TNFα during cancer immunity depend on context within the tumor microenvironment. Hence, in the presence of regulatory cell subsets, exhausted CD8^+^ and suppressive cytokine milieu, TNFα may promote tumor progression rather than immunity.

Beyond identifying the molecular axis, this work proposes a novel immunological concept of activation, where sequential signals provided in a time-dependent manner to T cells by different DCs at different anatomical sites govern T cell cytotoxicity. Importantly, this study also provides a roadmap to increase the therapeutic efficacy of DC vaccines and T cell immunotherapies.

## Supporting information

the file consists the supplemental figures and legends of the paper

## References

1. Hildner, K., Edelson, B.T., Purtha, W.E., Diamond, M., Matsushita, H., Kohyama, M., Calderon, B., Schraml, B.U., Unanue, E.R., and Diamond, M.S. (2008). Batf3 deficiency reveals a critical role for CD8α+ dendritic cells in cytotoxic T cell immunity. Science 322, 1097–1100.

2. Bhandarkar, V., Dinter, T., and Spranger, S. (2025). Architects of immunity: How dendritic cells shape CD8(+) T cell fate in cancer. Sci Immunol 10, eadf4726. 10.1126/sciimmunol.adf4726.

3. Mildner, A., and Jung, S. (2014). Development and function of dendritic cell subsets. Immunity 40, 642–656. 10.1016/j.immuni.2014.04.016 S1074-7613(14)00154-X [pii].

4. Merad, M., Sathe, P., Helft, J., Miller, J., and Mortha, A. (2013). The dendritic cell lineage: ontogeny and function of dendritic cells and their subsets in the steady state and the inflamed setting. Annual review of immunology 31, 563–604. 10.1146/annurev-immunol-020711074950.

5. Guilliams, M., Bruhns, P., Saeys, Y., Hammad, H., and Lambrecht, B.N. (2014). The function of Fcgamma receptors in dendritic cells and macrophages. Nat Rev Immunol 14, 94–108. 10.1038/nri3582 nri3582 [pii].

6. Bottcher, J.P., and Reis e Sousa, C. (2018). The Role of Type 1 Conventional Dendritic Cells in Cancer Immunity. Trends Cancer 4, 784–792. 10.1016/j.trecan.2018.09.001.

7. Guilliams, M., Dutertre, C.A., Scott, C.L., McGovern, N., Sichien, D., Chakarov, S., Van Gassen, S., Chen, J., Poidinger, M., De Prijck, S., et al. (2016). Unsupervised High-Dimensional Analysis Aligns Dendritic Cells across Tissues and Species. Immunity 45, 669–684. 10.1016/j.immuni.2016.08.015 S1074-7613(16)30339-9 [pii].

8. Salmon, H., Idoyaga, J., Rahman, A., Leboeuf, M., Remark, R., Jordan, S., Casanova-Acebes, M., Khudoynazarova, M., Agudo, J., Tung, N., et al. (2016). Expansion and Activation of CD103(+) Dendritic Cell Progenitors at the Tumor Site Enhances Tumor Responses to Therapeutic PD-L1 and BRAF Inhibition. Immunity 44, 924–938. 10.1016/j.immuni.2016.03.012.

9. Roberts, E.W., Broz, M.L., Binnewies, M., Headley, M.B., Nelson, A.E., Wolf, D.M., Kaisho, T., Bogunovic, D., Bhardwaj, N., and Krummel, M.F. (2016). Critical Role for CD103(+)/CD141(+) Dendritic Cells Bearing CCR7 for Tumor Antigen Trafficking and Priming of T Cell Immunity in Melanoma. Cancer Cell 30, 324–336. 10.1016/j.ccell.2016.06.003 S1535-6108(16)30263-X [pii].

10. Spranger, S., Dai, D., Horton, B., and Gajewski, T.F. (2017). Tumor-Residing Batf3 Dendritic Cells Are Required for Effector T Cell Trafficking and Adoptive T Cell Therapy. Cancer Cell 31, 711–723 e714. 10.1016/j.ccell.2017.04.003.

11. Spitzer, M.H., Carmi, Y., Reticker-Flynn, N.E., Kwek, S.S., Madhireddy, D., Martins, M.M., Gherardini, P.F., Prestwood, T.R., Chabon, J., Bendall, S.C., et al. (2017). Systemic Immunity Is Required for Effective Cancer Immunotherapy. Cell 168, 487–502 e415. S0092-8674(16)31738-X [pii] 10.1016/j.cell.2016.12.022.

12. Anguille, S., Smits, E.L., Lion, E., van Tendeloo, V.F., and Berneman, Z.N. (2014). Clinical use of dendritic cells for cancer therapy. Lancet Oncol 15, e257–267. 10.1016/S1470-2045(13)705850.

13. Wimmers, F., Schreibelt, G., Skold, A.E., Figdor, C.G., and De Vries, I.J. (2014). Paradigm Shift in Dendritic Cell-Based Immunotherapy: From in vitro Generated Monocyte-Derived DCs to Naturally Circulating DC Subsets. Front Immunol 5, 165. 10.3389/fimmu.2014.00165.

14. Palucka, K., and Banchereau, J. (2012). Cancer immunotherapy via dendritic cells. Nat Rev Cancer 12, 265–277. 10.1038/nrc3258.

15. Bol, K.F., Schreibelt, G., Gerritsen, W.R., de Vries, I.J., and Figdor, C.G. (2016). Dendritic CellBased Immunotherapy: State of the Art and Beyond. Clin Cancer Res 22, 1897–1906. 10.1158/1078-0432.CCR-15-1399.

16. Carreno, B.M., Magrini, V., Becker-Hapak, M., Kaabinejadian, S., Hundal, J., Petti, A.A., Ly, A., Lie, W.R., Hildebrand, W.H., Mardis, E.R., and Linette, G.P. (2015). Cancer immunotherapy. A dendritic cell vaccine increases the breadth and diversity of melanoma neoantigen-specific T cells. Science 348, 803–808. 10.1126/science.aaa3828.

17. Ott, P.A., Hu, Z., Keskin, D.B., Shukla, S.A., Sun, J., Bozym, D.J., Zhang, W., Luoma, A., GiobbieHurder, A., Peter, L., et al. (2017). An immunogenic personal neoantigen vaccine for patients with melanoma. Nature 547, 217–221. 10.1038/nature22991.

18. Sahin, U., Derhovanessian, E., Miller, M., Kloke, B.P., Simon, P., Lower, M., Bukur, V., Tadmor, A.D., Luxemburger, U., Schrors, B., et al. (2017). Personalized RNA mutanome vaccines mobilize poly-specific therapeutic immunity against cancer. Nature 547, 222–226. 10.1038/nature23003.

19. Butterfield, L.H. (2013). Dendritic cells in cancer immunotherapy clinical trials: are we making progress? Front Immunol 4, 454. 10.3389/fimmu.2013.00454.

20. Boudewijns, S., Bloemendal, M., de Haas, N., Westdorp, H., Bol, K.F., Schreibelt, G., Aarntzen, E., Lesterhuis, W.J., Gorris, M.A.J., Croockewit, A., et al. (2020). Autologous monocyte-derived DC vaccination combined with cisplatin in stage III and IV melanoma patients: a prospective, randomized phase 2 trial. Cancer Immunol Immunother 69, 477–488. 10.1007/s00262-01902466-x.

21. Yazdani, M., Jaafari, M.R., Verdi, J., Alani, B., Noureddini, M., and Badiee, A. (2020). Ex vivogenerated dendritic cell-based vaccines in melanoma: the role of nanoparticulate delivery systems. Immunotherapy 12, 333–349. 10.2217/imt-2019-0173.

22. Blass, E., Keskin, D.B., Tu, C.R., Forman, C., Vanasse, A., Sax, H.E., Shim, B., Chea, V., Kim, N., Carulli, I., et al. (2025). A multi-adjuvant personal neoantigen vaccine generates potent immunity in melanoma. Cell. 10.1016/j.cell.2025.06.019.

23. Gleisner, M.A., Pereda, C., Tittarelli, A., Navarrete, M., Fuentes, C., Avalos, I., Tempio, F., Araya, J.P., Becker, M.I., Gonzalez, F.E., et al. (2020). A heat-shocked melanoma cell lysate vaccine enhances tumor infiltration by prototypic effector T cells inhibiting tumor growth. J Immunother Cancer 8. 10.1136/jitc-2020-000999.

24. Bulgarelli, J., Tazzari, M., Granato, A.M., Ridolfi, L., Maiocchi, S., de Rosa, F., Petrini, M., Pancisi, E., Gentili, G., Vergani, B., et al. (2019). Dendritic Cell Vaccination in Metastatic Melanoma Turns “Non-T Cell Inflamed” Into “T-Cell Inflamed” Tumors. Front Immunol 10, 2353. 10.3389/fimmu.2019.02353.

25. Overwijk, W.W., Tsung, A., Irvine, K.R., Parkhurst, M.R., Goletz, T.J., Tsung, K., Carroll, M.W., Liu, C., Moss, B., Rosenberg, S.A., and Restifo, N.P. (1998). gp100/pmel 17 is a murine tumor rejection antigen: induction of “self”-reactive, tumoricidal T cells using high-affinity, altered peptide ligand. J Exp Med 188, 277–286. 10.1084/jem.188.2.277.

26. Carmi, Y., Spitzer, M.H., Linde, I.L., Burt, B.M., Prestwood, T.R., Perlman, N., Davidson, M.G., Kenkel, J.A., Segal, E., Pusapati, G.V., et al. (2015). Allogeneic IgG combined with dendritic cell stimuli induce antitumour T-cell immunity. Nature 521, 99–104. 10.1038/nature14424 nature14424 [pii].

27. Rafiq, K., Bergtold, A., and Clynes, R. (2002). Immune complex-mediated antigen presentation induces tumor immunity. The Journal of clinical investigation 110, 71–79. 10.1172/JCI15640.

28. Schuurhuis, D.H., van Montfoort, N., Ioan-Facsinay, A., Jiawan, R., Camps, M., Nouta, J., Melief, C.J., Verbeek, J.S., and Ossendorp, F. (2006). Immune complex-loaded dendritic cells are superior to soluble immune complexes as antitumor vaccine. J Immunol 176, 4573–4580.

29. Rahim, M.K., Okholm, T.L.H., Jones, K.B., McCarthy, E.E., Liu, C.C., Yee, J.L., Tamaki, S.J., Marquez, D.M., Tenvooren, I., Wai, K., et al. (2023). Dynamic CD8(+) T cell responses to cancer immunotherapy in human regional lymph nodes are disrupted in metastatic lymph nodes. Cell 186, 1127–1143 e1118. 10.1016/j.cell.2023.02.021.

30. Santana-Magal, N., Farhat-Younis, L., Gutwillig, A., Gleiberman, A., Rasoulouniriana, D., Tal, L., Netanely, D., Shamir, R., Blau, R., Feinmesser, M., et al. (2020). Melanoma-Secreted Lysosomes Trigger Monocyte-Derived Dendritic Cell Apoptosis and Limit Cancer Immunotherapy. Cancer Res 80, 1942–1956. 10.1158/0008-5472.CAN-19-2944.

31. Garris, C.S., Arlauckas, S.P., Kohler, R.H., Trefny, M.P., Garren, S., Piot, C., Engblom, C., Pfirschke, C., Siwicki, M., and Gungabeesoon, J. (2018). Successful anti-PD-1 cancer immunotherapy requires T cell-dendritic cell crosstalk involving the cytokines IFN-γ and IL-12. Immunity 49, 1148–1161. e1147.

32. Prokhnevska, N., Cardenas, M.A., Valanparambil, R.M., Sobierajska, E., Barwick, B.G., Jansen, C., Reyes Moon, A., Gregorova, P., delBalzo, L., Greenwald, R., et al. (2023). CD8(+) T cell activation in cancer comprises an initial activation phase in lymph nodes followed by effector differentiation within the tumor. Immunity 56, 107–124 e105. 10.1016/j.immuni.2022.12.002.

33. Sheinin, R., Salomon, K., Yeini, E., Dulberg, S., Kaminitz, A., Satchi-Fainaro, R., Sharan, R., and Madi, A. (2024). interFLOW: maximum flow framework for the identification of factors mediating the signalling convergence of multiple receptors. NPJ Syst Biol Appl 10, 66. 10.1038/s41540-024-00391-z.

34. Marciscano, A.E., and Anandasabapathy, N. (2021). The role of dendritic cells in cancer and anti-tumor immunity. Semin Immunol 52, 101481. 10.1016/j.smim.2021.101481.

35. Matzinger, P. (1994). Tolerance, danger, and the extended family. Annual review of immunology 12, 991–1045. 10.1146/annurev.iy.12.040194.005015.

36. Janeway, C.A., Jr., and Medzhitov, R. (2002). Innate immune recognition. Annual review of immunology 20, 197–216. 10.1146/annurev.immunol.20.083001.084359.

37. Bennett, S.R., Carbone, F.R., Karamalis, F., Flavell, R.A., Miller, J.F., and Heath, W.R. (1998). Help for cytotoxic-T-cell responses is mediated by CD40 signalling. Nature 393, 478–480. 10.1038/30996.

38. Schoenberger, S.P., Toes, R.E., van der Voort, E.I., Offringa, R., and Melief, C.J. (1998). T-cell help for cytotoxic T lymphocytes is mediated by CD40-CD40L interactions. Nature 393, 480483. 10.1038/31002.

39. Joffre, O.P., Segura, E., Savina, A., and Amigorena, S. (2012). Cross-presentation by dendritic cells. Nat Rev Immunol 12, 557–569. 10.1038/nri3254.

40. Schlitzer, A., and Ginhoux, F. (2014). Organization of the mouse and human DC network. Current opinion in immunology 26, 90–99.

41. Martín-Fontecha, A., Sebastiani, S., Höpken, U.E., Uguccioni, M., Lipp, M., Lanzavecchia, A., and Sallusto, F. (2003). Regulation of dendritic cell migration to the draining lymph node: impact on T lymphocyte traffic and priming. The Journal of experimental medicine 198, 615621.

42. Roberts, E.W., Broz, M.L., Binnewies, M., Headley, M.B., Nelson, A.E., Wolf, D.M., Kaisho, T., Bogunovic, D., Bhardwaj, N., and Krummel, M.F. (2016). Critical Role for CD103(+)/CD141(+) Dendritic Cells Bearing CCR7 for Tumor Antigen Trafficking and Priming of T Cell Immunity in Melanoma. Cancer Cell 30, 324–336. 10.1016/j.ccell.2016.06.003.

43. Allan, R.S., Waithman, J., Bedoui, S., Jones, C.M., Villadangos, J.A., Zhan, Y., Lew, A.M., Shortman, K., Heath, W.R., and Carbone, F.R. (2006). Migratory dendritic cells transfer antigen to a lymph node-resident dendritic cell population for efficient CTL priming. Immunity 25, 153162. 10.1016/j.immuni.2006.04.017.

44. Reste, M., Ajazi, K., Sayi-Yazgan, A., Jankovic, R., Bufan, B., Brandau, S., Baekkevold, E.S., Petitprez, F., Lindstedt, M., Adema, G.J., and Almeida, C.R. (2024). The role of dendritic cells in tertiary lymphoid structures: implications in cancer and autoimmune diseases. Front Immunol 15, 1439413. 10.3389/fimmu.2024.1439413.

45. Spranger, S., Koblish, H.K., Horton, B., Scherle, P.A., Newton, R., and Gajewski, T.F. (2014). Mechanism of tumor rejection with doublets of CTLA-4, PD-1/PD-L1, or IDO blockade involves restored IL-2 production and proliferation of CD8+ T cells directly within the tumor microenvironment. Journal for immunotherapy of cancer 2, 1–14.

46. de Mingo Pulido, Á., Gardner, A., Hiebler, S., Soliman, H., Rugo, H.S., Krummel, M.F., Coussens, L.M., and Ruffell, B. (2018). TIM-3 regulates CD103+ dendritic cell function and response to chemotherapy in breast cancer. Cancer cell 33, 60–74. e66.

47. Ruffell, B., Chang-Strachan, D., Chan, V., Rosenbusch, A., Ho, C.M., Pryer, N., Daniel, D., Hwang, E.S., Rugo, H.S., and Coussens, L.M. (2014). Macrophage IL-10 blocks CD8+ T celldependent responses to chemotherapy by suppressing IL-12 expression in intratumoral dendritic cells. Cancer Cell 26, 623–637. 10.1016/j.ccell.2014.09.006.

48. Aarntzen, E.H., Schreibelt, G., Bol, K., Lesterhuis, W.J., Croockewit, A.J., de Wilt, J.H., van Rossum, M.M., Blokx, W.A., Jacobs, J.F., Duiveman-de Boer, T., et al. (2012). Vaccination with mRNA-electroporated dendritic cells induces robust tumor antigen-specific CD4+ and CD8+ T cells responses in stage III and IV melanoma patients. Clin Cancer Res 18, 5460–5470. 10.1158/1078-0432.CCR-11-3368.

49. De Keersmaecker, B., Claerhout, S., Carrasco, J., Bar, I., Corthals, J., Wilgenhof, S., Neyns, B., and Thielemans, K. (2020). TriMix and tumor antigen mRNA electroporated dendritic cell vaccination plus ipilimumab: link between T-cell activation and clinical responses in advanced melanoma. J Immunother Cancer 8. 10.1136/jitc-2019-000329.

50. Bentzen, A.K., Such, L., Jensen, K.K., Marquard, A.M., Jessen, L.E., Miller, N.J., Church, C.D., Lyngaa, R., Koelle, D.M., Becker, J.C., et al. (2018). T cell receptor fingerprinting enables indepth characterization of the interactions governing recognition of peptide-MHC complexes. Nat Biotechnol. 10.1038/nbt.4303.

51. Birnbaum, M.E., Mendoza, J.L., Sethi, D.K., Dong, S., Glanville, J., Dobbins, J., Ozkan, E., Davis, M.M., Wucherpfennig, K.W., and Garcia, K.C. (2014). Deconstructing the peptide-MHC specificity of T cell recognition. Cell 157, 1073–1087. 10.1016/j.cell.2014.03.047.

52. Kuhn, S., Hyde, E.J., Yang, J., Rich, F.J., Harper, J.L., Kirman, J.R., and Ronchese, F. (2013). Increased numbers of monocyte-derived dendritic cells during successful tumor immunotherapy with immune-activating agents. J Immunol 191, 1984–1992. 10.4049/jimmunol.1301135.

53. Kuhn, S., Yang, J., and Ronchese, F. (2015). Monocyte-Derived Dendritic Cells Are Essential for CD8(+) T Cell Activation and Antitumor Responses After Local Immunotherapy. Front Immunol 6, 584. 10.3389/fimmu.2015.00584.

54. Serbina, N.V., Salazar-Mather, T.P., Biron, C.A., Kuziel, W.A., and Pamer, E.G. (2003). TNF/iNOS-producing dendritic cells mediate innate immune defense against bacterial infection. Immunity 19, 59–70. 10.1016/s1074-7613(03)00171-7.

55. Fei, M., Bhatia, S., Oriss, T.B., Yarlagadda, M., Khare, A., Akira, S., Saijo, S., Iwakura, Y., Fallert Junecko, B.A., Reinhart, T.A., et al. (2011). TNF-alpha from inflammatory dendritic cells (DCs) regulates lung IL-17A/IL-5 levels and neutrophilia versus eosinophilia during persistent fungal infection. Proc Natl Acad Sci U S A 108, 5360–5365. 10.1073/pnas.1015476108.

56. Bosschaerts, T., Guilliams, M., Stijlemans, B., Morias, Y., Engel, D., Tacke, F., Herin, M., De Baetselier, P., and Beschin, A. (2010). Tip-DC development during parasitic infection is regulated by IL-10 and requires CCL2/CCR2, IFN-gamma and MyD88 signalling. PLoS Pathog 6, e1001045. 10.1371/journal.ppat.1001045.

57. Lowes, M.A., Chamian, F., Abello, M.V., Fuentes-Duculan, J., Lin, S.L., Nussbaum, R., Novitskaya, I., Carbonaro, H., Cardinale, I., Kikuchi, T., et al. (2005). Increase in TNF-alpha and inducible nitric oxide synthase-expressing dendritic cells in psoriasis and reduction with efalizumab (anti-CD11a). Proc Natl Acad Sci USA 102, 19057–19062. 10.1073/pnas.0509736102.

58. Marigo, I., Zilio, S., Desantis, G., Mlecnik, B., Agnellini, A.H.R., Ugel, S., Sasso, M.S., Qualls, J.E., Kratochvill, F., Zanovello, P., et al. (2016). T Cell Cancer Therapy Requires CD40-CD40L Activation of Tumor Necrosis Factor and Inducible Nitric-Oxide-Synthase-Producing Dendritic Cells. Cancer Cell 30, 377–390. 10.1016/j.ccell.2016.08.004.

59. Carswell, E.A., Old, L.J., Kassel, R.L., Green, S., Fiore, N., and Williamson, B. (1975). An endotoxin-induced serum factor that causes necrosis of tumors. Proc Natl Acad Sci U S A 72, 3666–3670. 10.1073/pnas.72.9.3666.

60. Dadiani, M., Necula, D., Kahana-Edwin, S., Oren, N., Baram, T., Marin, I., Morzaev-Sulzbach, D., Pavlovski, A., Balint-Lahat, N., Anafi, L., et al. (2020). TNFR2+ TILs are significantly associated with improved survival in triple-negative breast cancer patients. Cancer Immunol Immunother 69, 1315–1326. 10.1007/s00262-020-02549-0.

61. Tam, E.M., Fulton, R.B., Sampson, J.F., Muda, M., Camblin, A., Richards, J., Koshkaryev, A., Tang, J., Kurella, V., Jiao, Y., et al. (2019). Antibody-mediated targeting of TNFR2 activates CD8(+) T cells in mice and promotes antitumor immunity. Sci Transl Med 11. 10.1126/scitranslmed.aax0720.

62. Jones, A.L., O’Brien, M.E., Lorentzos, A., Viner, C., Hanrahan, A., Moore, J., Millar, J.L., and Gore, M.E. (1992). A randomised phase II study of carmustine alone or in combination with tumour necrosis factor in patients with advanced melanoma. Cancer Chemother Pharmacol 30, 73–76. 10.1007/BF00686489.

63. Roberts, N.J., Zhou, S., Diaz, L.A., Jr., and Holdhoff, M. (2011). Systemic use of tumor necrosis factor alpha as an anticancer agent. Oncotarget 2, 739–751. 10.18632/oncotarget.344.

64. Moore, R.J., Owens, D.M., Stamp, G., Arnott, C., Burke, F., East, N., Holdsworth, H., Turner, L., Rollins, B., Pasparakis, M., et al. (1999). Mice deficient in tumor necrosis factor-alpha are resistant to skin carcinogenesis. Nat Med 5, 828–831. 10.1038/10552.

65. Arnott, C.H., Scott, K.A., Moore, R.J., Robinson, S.C., Thompson, R.G., and Balkwill, F.R. (2004). Expression of both TNF-alpha receptor subtypes is essential for optimal skin tumour development. Oncogene 23, 1902–1910. 10.1038/sj.onc.1207317.

66. Montfort, A., Colacios, C., Levade, T., Andrieu-Abadie, N., Meyer, N., and Segui, B. (2019). The TNF Paradox in Cancer Progression and Immunotherapy. Front Immunol 10, 1818. 10.3389/fimmu.2019.01818.

