## Supplementary material for "Successful dendritic cell vaccines require lasting in-situ TNFa secretion to license antitumor CD8^+^ T cell cytotoxicity": the file consists the supplemental figures and legends of the paper

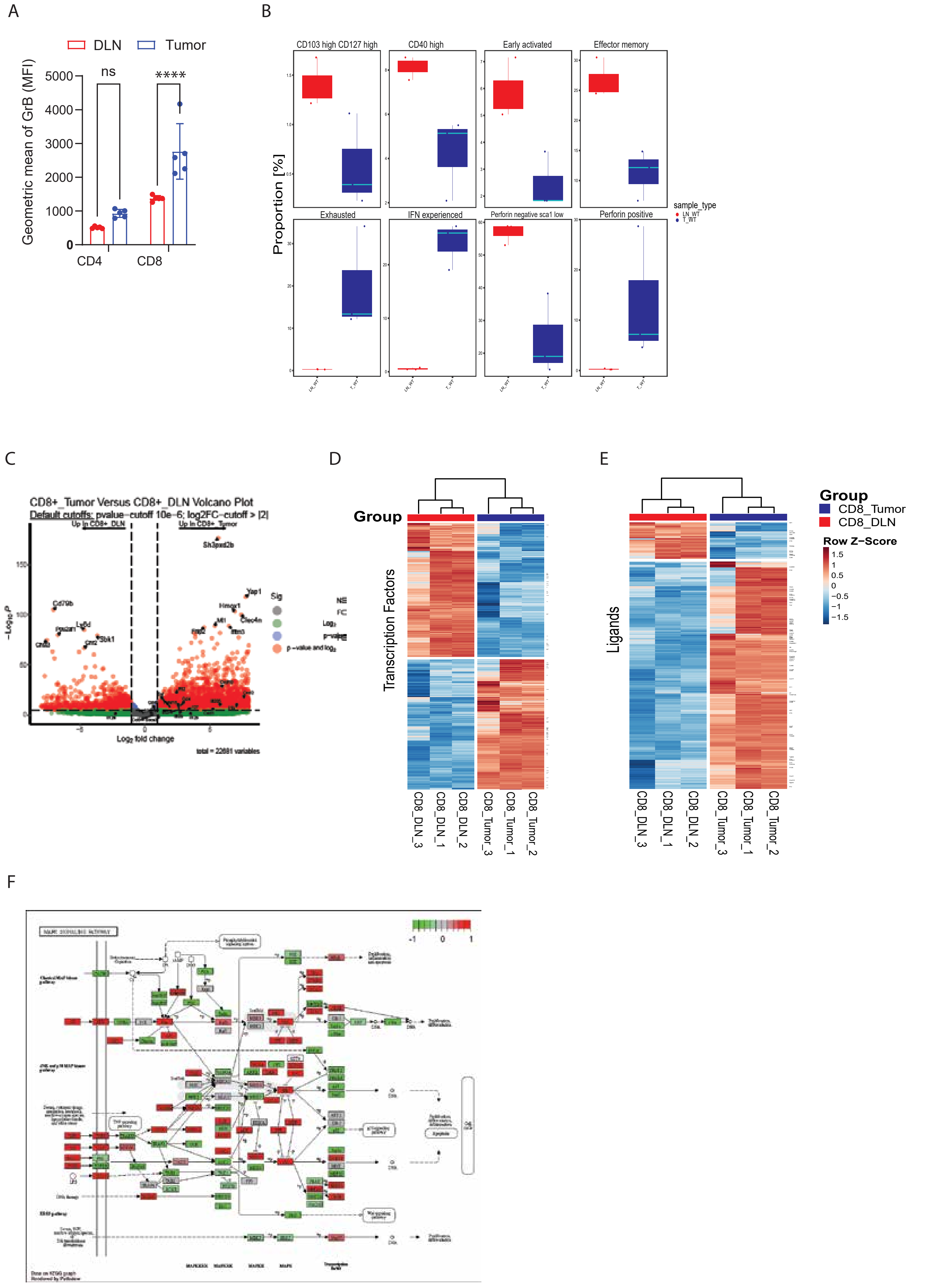
A
DLN
Tumor
Geometric mean of GrB (MFI)
ns
****
5000
4000
3000
2000
1000
0
CD4
CD8
B
CD103 high CD127 high
CD40 high
Early activated
Effector memory
Exhausted
IFN experienced
Perforin negative sca1 low
Perforin positive
Proportion [%]
sample_type
LN_WT
T_WT
C
CD8+_Tumor Versus CD8+_DLN Volcano Plot
Default cutoffs: pvalue-cutoff 10e-6; log2FC-cutoff > |2|
Up in CD8+_DLN
Up in CD8+_Tumor
Sh3pxd2b
Yap1
Cd79b
Sig
NS
FC
p−value and log
Log fold change
total = 22681 variables
D
Group
Transcription Factors
CD8_DLN_3
CD8_DLN_1
CD8_DLN_2
CD8_Tumor_3
CD8_Tumor_1
CD8_Tumor_2
E
Ligands
Group
CD8_Tumor
CD8_DLN
Row Z−Score
1.5
1
0.5
0
−0.5
−1
−1.5
F
MAPK SIGNALING PATHWAY
-1
0
1

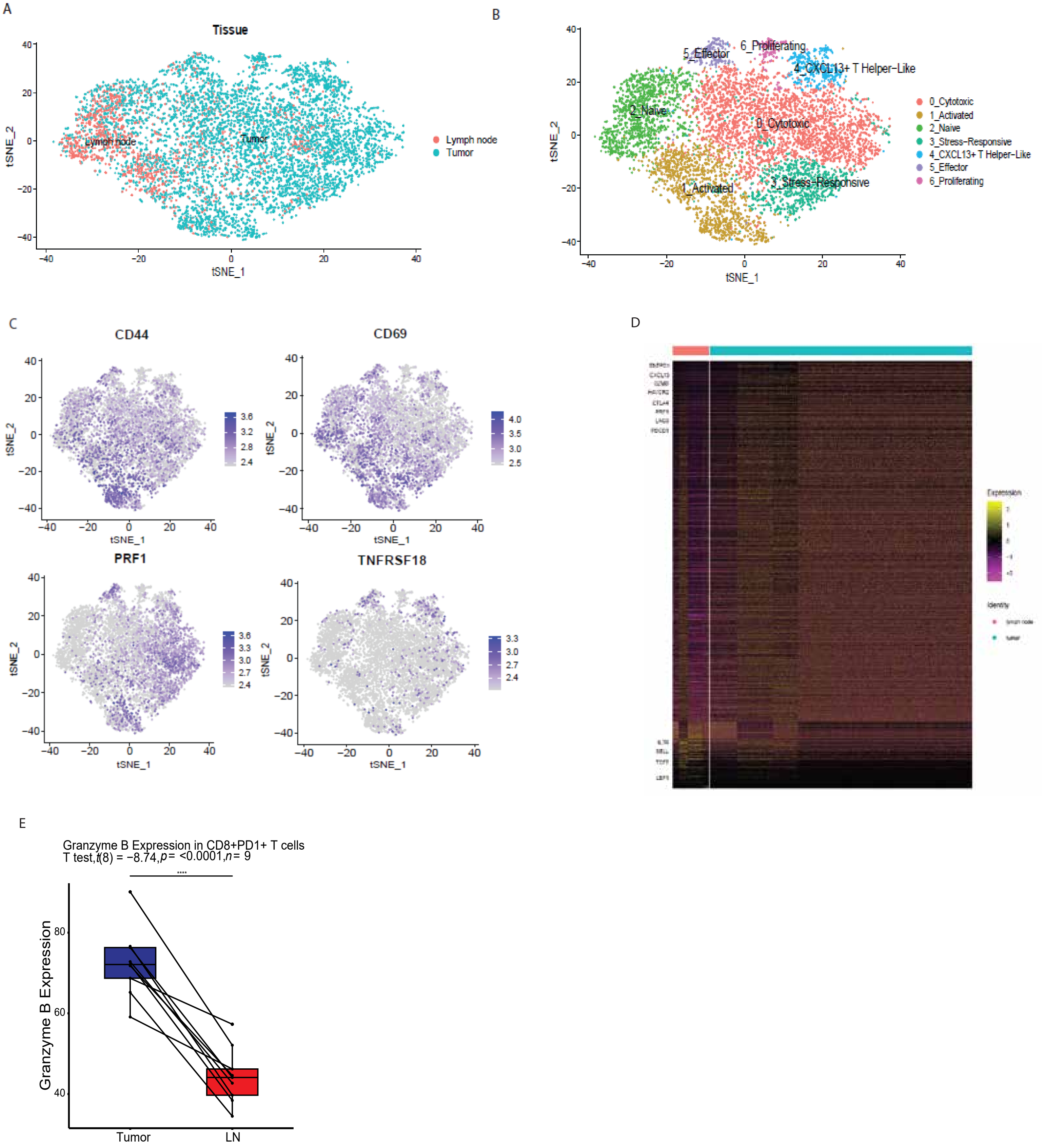
A
Tissue
Lymph node
Tumor
tSNE_1
tSNE_2
B
0_Cytotoxic
1_Activated
2_Naive
3_Stress-Responsive
4_CXCL13+ T Helper-Like
5_Effector
6_Proliferating
C
CD44
CD69
PRF1
TNFRSF18
D
E
Granzyme B Expression in CD8+PD1+ T cells
T test, t(8) = −8.74, p = <0.0001, n = 9
Granzyme B Expression
Tumor
LN

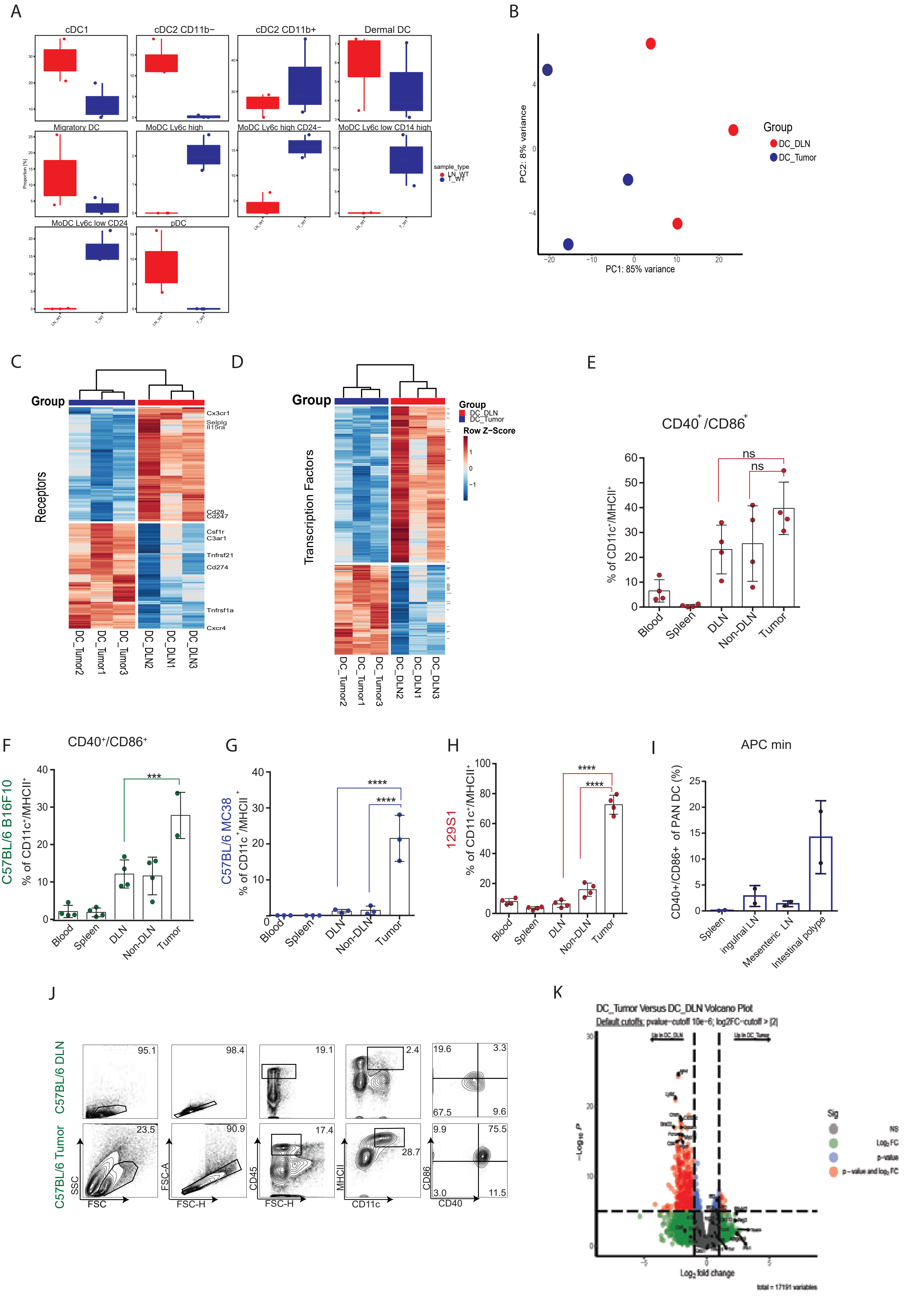
A
cDC1
cDC2 CD11b−
cDC2 CD11b+
Dermal DC
Migratory DC
MoDC Ly6c high
MoDC Ly6c high CD24−
MoDC Ly6c low CD14 high
MoDC Ly6c low CD24
pDC
Proportion [%]
sample_type
LN_WT
T_WT
B
PC2: 8% variance
PC1: 85% variance
Group
DC_DLN
DC_Tumor
C
Group
Receptors
Cx3cr1
Selplg
Il15ra
Cd28
Cd247
Csf1r
C3ar1
Tnfrsf21
Cd274
Tnfrsf1a
Cxcr4
DC_Tumor2
DC_Tumor1
DC_Tumor3
DC_DLN2
DC_DLN1
DC_DLN3
D
Group
Transcription Factors
Row Z−Score
E
CD40+/CD86+
% of CD11c+/MHCII+
ns
Blood
Spleen
DLN
Non-DLN
Tumor
F
CD40+/CD86+
C57BL/6 B16F10
% of CD11c+/MHCII+
***
G
C57BL/6 MC38
% of CD11c+/MHCII+
****
H
129S1
% of CD11c+/MHCII+
****
I
APC min
CD40+/CD86+ of PAN DC (%)
Spleen
ingulnal LN
Mesenteric LN
Intestinal polype
J
C57BL/6 DLN
C57BL/6 Tumor
SSC
FSC
FSC-A
FSC-H
CD45
MHCII
CD11c
CD86
CD40
95.1
98.4
19.1
2.4
19.6
3.3
67.5
9.6
23.5
90.9
17.4
28.7
9.9
75.5
3.0
11.5
K
DC_Tumor Versus DC_DLN Volcano Plot
Default cutoffs: pvalue-cutoff 10e-6; log2FC-cutoff > |2|
Log2 fold change
Sig
NS
Log2 FC
p-value
p-value and log2 FC
total = 17191 variables

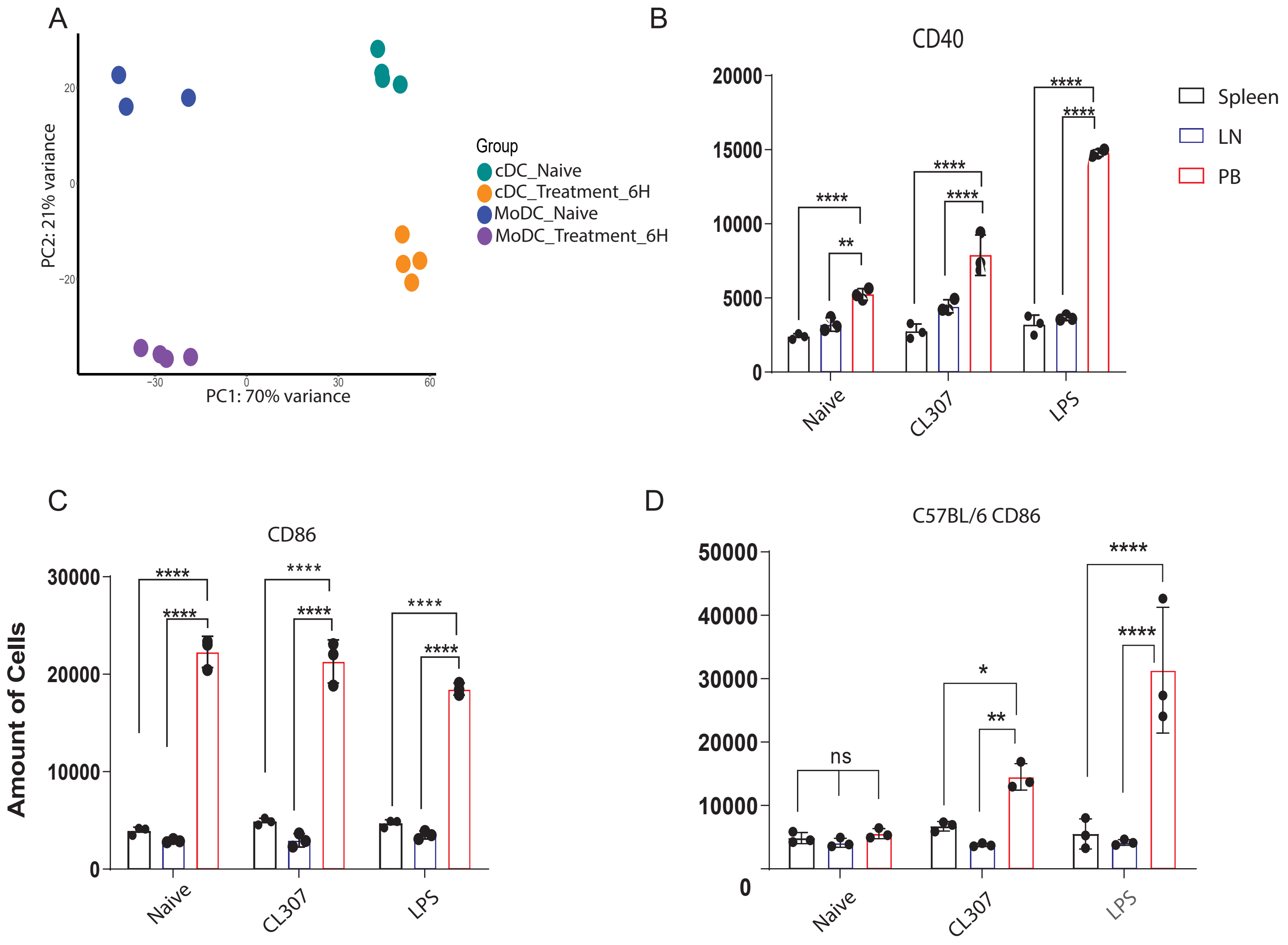
A
PC2: 21% variance
PC1: 70% variance
Group
cDC_Naive
cDC_Treatment_6H
MoDC_Naive
MoDC_Treatment_6H
B
CD40
Spleen
LN
PB
Naive
CL307
LPS
C
CD86
Amount of Cells
D
C57BL/6 CD86
ns

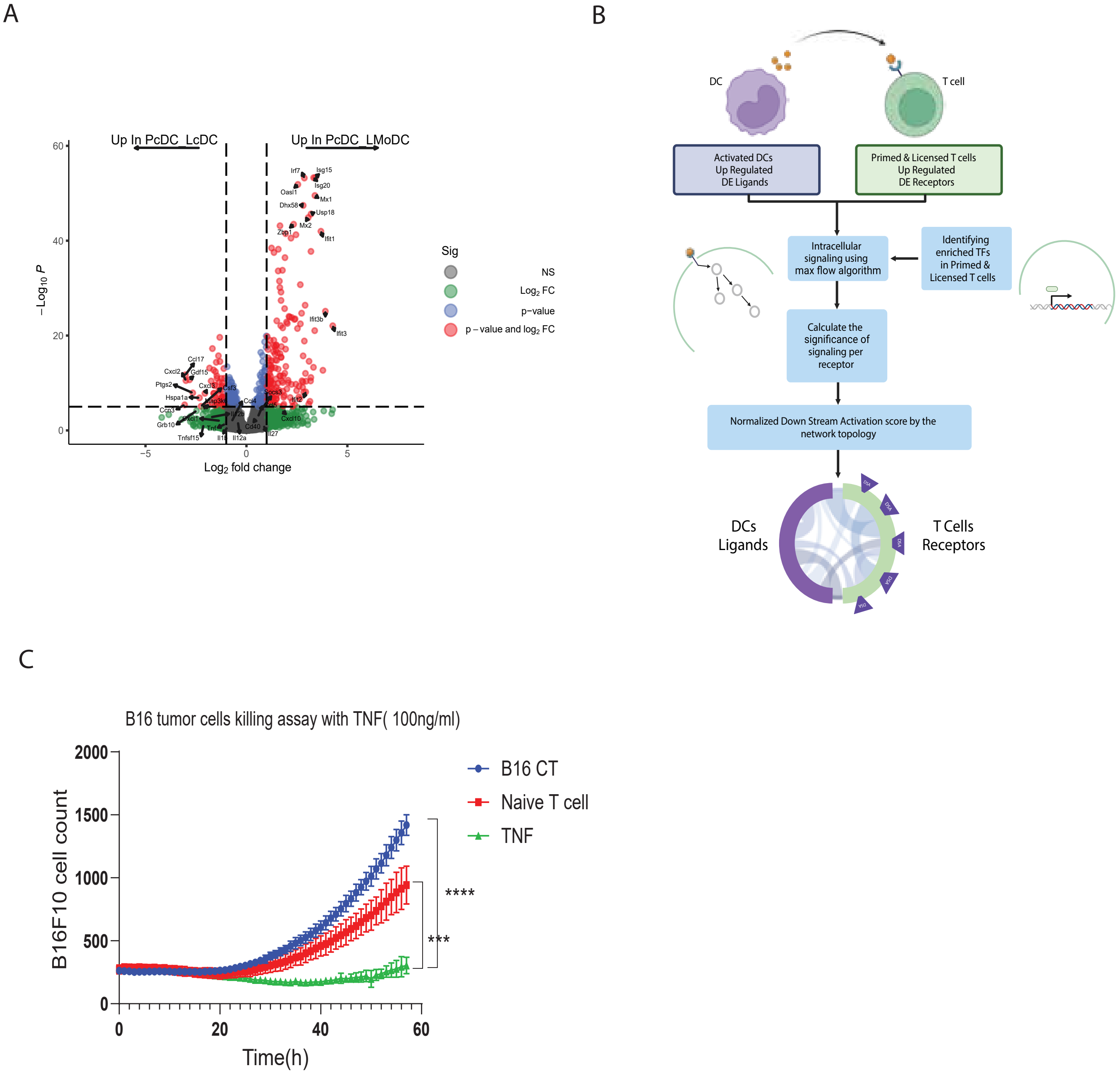

A
Up In PcDC_LcDC
Up In PcDC_LMoDC
−Log10 P
Log2 fold change
Sig
NS
Log2 FC
p−value
p − value and log2 FC
B
DC
T cell
Activated DCs
Up Regulated
DE Ligands
Primed & Licensed T cells
Up Regulated
DE Receptors
Intracellular
signaling using
max flow algorithm
Identifying
enriched TFs
in Primed &
Licensed T cells
Calculate the
significance of
signaling per
receptor
Normalized Down Stream Activation score by the
network topology
DCs
Ligands
T Cells
Receptors
C
B16 tumor cells killing assay with TNF( 100ng/ml)
B16F10 cell count
Time(h)
B16 CT
Naive T cell
TNF
****
***

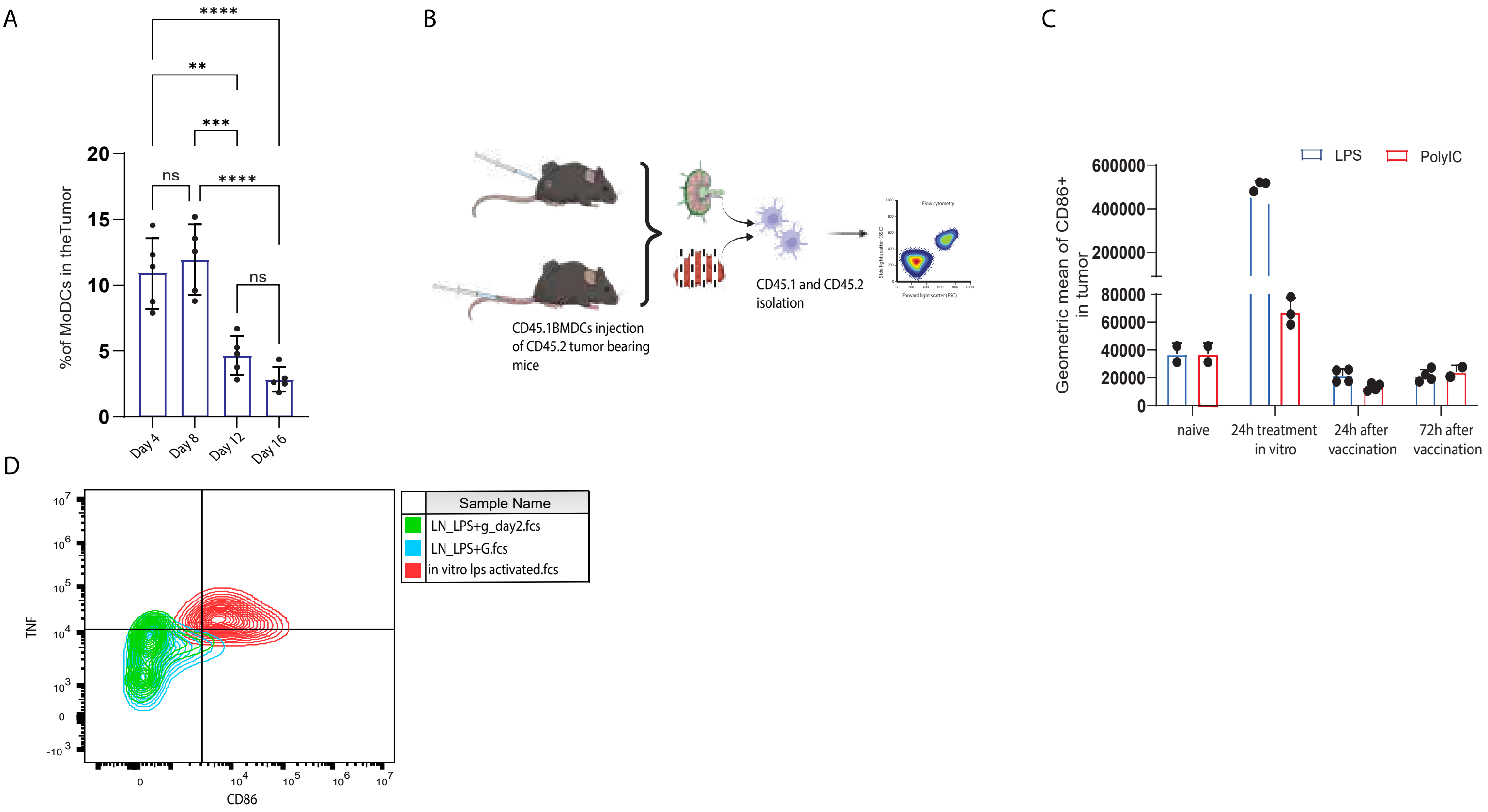
A
****
**
***
ns
****
ns
%of MoDCs in the Tumor
20
15
10
5
0
Day 4
Day 8
Day 12
Day 16
B
CD45.1BMDCs injection of CD45.2 tumor bearing mice
CD45.1 and CD45.2 isolation
Flow cytometry
C
LPS
PolyIC
Geometric mean of CD86+ in tumor
600000
400000
200000
80000
60000
40000
20000
0
naive
24h treatment in vitro
24h after vaccination
72h after vaccination
D
Sample Name
LN_LPS+g_day2.fcs
LN_LPS+G.fcs
in vitro lps activated.fcs
TNF
CD86

**Supplemental Figure Legends**

*Figure S1: Murine $CD8^+$ T cells from the tumor and DLN manifest different activation phenotype***.** (A) Mean fluorescence intensity (MFI) of Granzyme B expression in CD8 isolated from DLN ant tumors of day 10 B16F10 tumor bearing mice (n=4). (B) Proportion of $CD8^+$ T cells in each cluster in mouse tumor and DLN (n=3). (C) Volcano plot showing upregulated differential expression of proteins between $CD8^+$ T cells isolated from DLN and tumors (n=3). (D-E) Hierarchal cluster of significantly changed genes in $CD8^+$ T cells isolated from DLN and tumors from mice bearing day 10 B16F10 (n=3) (D) transcriptional factors, (E) ligands. (F) KEGG analyses on differentially expressed MAPK signalling pathway genes in T cells from DLN vs. Tumor (n=3).

*Figure S2: Human $CD8^+$ T cells from the tumor and DLN manifest different activation phenotype***.** (A) t-SNE plot illustrating the tissue origin (tumor or lymph node) of 7,987 $CD8^+$ T cells isolated from tumor and lymph node samples. (B) t-SNE plot showing the transcriptional profiles of 7,987 $CD8^+$ T cells isolated from tumor and lymph node samples. Each point represents an individual cell, colored and annotated by cluster identity. (C) Feature plots demonstrating the expression levels of selected top DE genes across individual cells, with a minimum expression cut-off set at the 25$^{th}$ quantile. (D) Heatmap displaying the transcriptional differences between tumor-derived and LN-derived effector $CD8^+$ T cells, comparing differentially expressed genes. Yellow indicates upregulation, while purple indicates downregulation. (E) Granzyme B expression in $CD8^+PDL1^+$ T cells from paired tumor and LN samples. Each line connects paired values from individual samples, illustrating differences in expression between tumor and LN tissues (paired t-test, $t$ (8) = -8.74, $p < 0.0001$, $n$ = 9).

*Figure S3: MoDC manifest inflammatory phenotype across different mouse models.* (A) Proportion of DC in each cluster in mouse Tumor and DLN (n=3). (B) Principal component analysis of gene expression patterns comparing tumor and DLN DCs from tumor bearing C57BL/6 mice (n=3). (C-F) Percentage of cd11c+/MHCII + cells expressing activation markers CD86/CD40 isolated from either blood, spleen, DLN, N-DLN and tumor of (E-F) C75BL/6 tumor bearing mice (E) 129S1 tumor bearing mice (n=4) (F). (G) Percentage of CD86/CD40 expression on $CD11c^+$ MHCII high cells isolated from either Spleen, Inguinal LN, Mesenteric LN, Intestinal polyp (n=4). (H) FACS analysis of DC expressing CD86/CD40 isolated from DLN and tumor of B16F10 tumor-bearing C75BL/6 mice (day 10; n=4). (I-J) Hierarchal cluster of significantly changed genes (J) Receptors, (K) Transcriptional factors. in DCs cells isolated from DLN vs Tumor of tumor bearing C57BL/6 mice (n=3). (K) Volcano plot showing upregulated differential expression of proteins between DCs cells isolated from either DLN or Tumor (n=3).

Figure S4: *MoDC express higher levels of costimulatory molecules compared to cDC.* (A) Principal component analysis of gene expression patterns comparing naïve vs 6h LPS activated cDCs or MoDCs (n=3). (B) Geometric mean of CD40 expressed in naïve, CL307 or LPS activated DCs isolated from either peripheral blood, LN or spleen (n=3). (C) Geometric mean of CD86 expressed in naïve, CL307 or LPS activated DCs isolated from either peripheral blood, LN or spleen (n=3). (D) Amount of CD86+ cells expressed in naïve, CL307 or LPS activated DCs isolated from either peripheral blood, LN or spleen of C57BL/6 mice (n=3).

*Figure S5: TNF□ elicit T cell cytotoxicity*. (A) Volcano plot showing differential expression of proteins between primed with cDCs and licensed with cDCs vs MoDCs $CD8^+$ T cells, Default cutoffs: p-value−cutoff 10e−6; log2FC−cutoff > |2|, total = 11399 variables (n=3). (B) Workflow illustrating the process developed to construct the ligand-receptor interaction plot and calculate DSA values using the algorithm. (C) Mean B16F10 counts over time following incubation with primed $CD8^+$ T cells with or without TNF (n=6).

*Figure S6: DC rapidly loses their activation phenotype.* (A) Mean percentage of MoDCs cells isolated from tumor of tumor bearing mice on different days (n=5). (B) Illustration of experimental outline (https://biorender.com/). (C) Mean percentage of CD86 expression on $CD45.1^+$ MoDCs detected in B16F10 tumors at 24- and 72-hours postvaccination. MoDCs were activated overnight with either LPS or Poly I:C, pulsed with gp100 peptide, and injected s.c. into CD45.2 mice 10 days after B16F10 tumor challenge (n=2,3,4 or 5). (D) FACS analysis of DCs expressing TNF, in vitro and CD45.1 DCs isolated from the LN of B16F10 tumor-bearing C75BL/6 mice (day 10/11; *n* = 5).
